# A Structural Mechanism for Noncanonical GPCR Signal Transduction in the Hedgehog Pathway

**DOI:** 10.1101/2024.10.31.621410

**Authors:** William P. Steiner, Nathan Iverson, Guibing Liu, Varun Venkatakrishnan, Jian Wu, Tomasz Maciej Stepniewski, Zachary Michaelson, Jan W. Bröckel, Ju-Fen Zhu, Jessica Bruystens, Annabel Lee, Isaac Nelson, Daniela Bertinetti, Corvin D. Arveseth, Gerald Tan, Paul Spaltenstein, Jiewei Xu, Ruth Hüttenhain, Michael Kay, Friedrich W. Herberg, Erhu Cao, Jana Selent, Ganesh S. Anand, Roland L. Dunbrack, Susan S. Taylor, Benjamin R. Myers

## Abstract

The Hedgehog (Hh) signaling pathway is fundamental to embryogenesis, tissue homeostasis, and cancer. Hh signals are transduced via an unusual mechanism: upon agonist-induced phosphorylation, the noncanonical G protein-coupled receptor SMOOTHENED (SMO) binds the catalytic subunit of protein kinase A (PKA-C) and physically blocks its enzymatic activity. By combining computational structural approaches with biochemical and functional studies, we show that SMO mimics strategies prevalent in canonical GPCR and PKA signaling complexes, despite little sequence or secondary structural homology. An intrinsically disordered region of SMO binds the PKA-C active site, resembling the PKA regulatory subunit (PKA-R) / PKA-C holoenzyme, while the SMO transmembrane domain binds a conserved PKA-C interaction hub, similar to other GPCR-effector complexes. In contrast with prevailing GPCR signal transduction models, phosphorylation of SMO promotes intramolecular electrostatic interactions that stabilize key structural elements within the SMO cytoplasmic domain, thereby remodeling it into a PKA-inhibiting conformation. Our work provides a structural mechanism for a central step in the Hh cascade and defines a paradigm for disordered GPCR domains to transmit signals intracellularly.

## INTRODUCTION

Communication between GPCRs and PKA is fundamental to human physiology, and malfunctions in GPCR-PKA communication can lead to disease^1,2^. Canonical GPCRs control PKA activity by coupling to heterotrimeric G proteins which regulate formation of cyclic AMP, a second messenger that unleashes PKA-C subunits from inhibition by PKA-R subunits in PKA holoenzymes^1,2^. In contrast, during vertebrate Hh signal transduction in the primary cilium^3–5^, the noncanonical GPCR SMO is activated by sterols^6–12^, which triggers insertion of a PKA pseudosubstrate motif within the SMO intracellular membrane-proximal C-terminus (pCT) into the PKA-C active site^13–15^. Binding of this pseudosubstrate motif sequesters PKA-C at the membrane and interrupts its catalytic cycle^13–15^. Consequently, glioma-associated (GLI) transcription factors are relieved from phosphorylation-induced inhibition, enabling expression of Hh pathway target genes that mediate cell proliferation or differentiation^13–15^.

SMO / PKA-C interactions are regulated such that only the active state of SMO can inhibit PKA-C, ensuring appropriate regulation of GLI transcriptional activity and preventing pathological outcomes^13,15^. Thus, GPCR kinases 2 and 3 (GRK2/3) recognize SMO specifically in its active, sterol-bound state and phosphorylate SMO at sites dispersed throughout its pCT, thereby enhancing interactions between the SMO pseudosubstrate motif and PKA-C^13,15^. Although phosphorylated GPCRs often bind cytoplasmic signaling proteins indirectly via β-arrestin scaffolds^16–19^, *in vitro* reconstitution studies with purified proteins established that GRK2/3 phosphorylation of SMO triggers a direct SMO / PKA-C interaction^15^. The SMO-GRK2/3-PKA pathway is essential to Hh signal transduction^13–15,20–23^, but how SMO binds PKA-C in structural terms, and how GRK2/3 phosphorylation facilitates this process, are unknown.

Numerous structural studies have explored how canonical GPCRs bind conventional downstream effectors^16–18,24^, as well as how PKA-C engages classical pseudosubstrates^2,25–27^. The SMO / PKA-C interaction is distinct from these signaling complexes in two fundamental ways. First, PKA-C is unique among GPCR effectors: it is unrelated to G proteins or β-arrestins and contains no obvious GPCR-interacting domains, such as the positively charged phosphopeptide-binding groove that enables β-arrestins to selectively interact with the C-termini of active, phosphorylated GPCRs^16,17^. Second, unlike conventional PKA inhibitors, such as the heat-stable protein kinase inhibitor (PKI) proteins or PKA-R subunits, which interact with PKA-C constitutively or under cAMP-depleted conditions, respectively^25,28^, SMO requires GRK2/3 phosphorylation to bind and inhibit PKA-C during Hh signal transduction^13–15^. Consistent with this unprecedented mode of PKA regulation, SMO only displays homology to PKI proteins and PKA-R subunits within its pseudosubstrate region^14^, and does not appear to encode any of the additional sequence or secondary structural elements in these proteins that are needed for efficient PKA-C binding^29–32^. Thus, existing GPCR and PKA structures offer little insight into the SMO / PKA-C complex.

The intracellular segments of both canonical and noncanonical GPCRs (including SMO) are predicted to be intrinsically disordered regions (IDRs)^33^. IDRs adopt a continuum of conformational states^34–37^, presenting a formidable challenge for biophysical and structural characterization. Indeed, these GPCR IDRs – critical for signaling – are largely unresolved in or entirely absent from most GPCR crystallography or cryo-electron microscopy structures^33,38,39^, including the >20 SMO structures solved to date^7,11,40–43^. To overcome the limitations of traditional approaches, we analyzed the SMO / PKA-C complex by integrating computational structure prediction and molecular dynamics (MD) simulations with biochemical, biophysical, and functional studies. The resulting structural model reveals key mechanistic insights into this critical but previously inaccessible signaling complex.

## RESULTS

### AlphaFold modeling of the SMO / PKA-C complex

Unlike the well-folded SMO extracellular and 7-transmembrane (7TM) domains, which have been extensively studied via empirical structural methods^7,11,40–43^, the SMO pCT remains structurally uncharacterized. Prior computational predictions suggest that the SMO pCT is an IDR^13^, consistent with the inability to resolve this region of SMO in cryo-electron microscopy^42^. To determine whether the SMO pCT is an IDR, we analyzed a soluble portion of SMO encompassing almost the entire SMO pCT (residues 565-657) via circular dichroism (CD) spectroscopy. The purified SMO pCT showed a mixed weak ɑ-helical and random coil fingerprint **(Extended data Fig. 1a**), indicating limited secondary structure or an ensemble of disordered and partially structured states.

**Figure 1:**
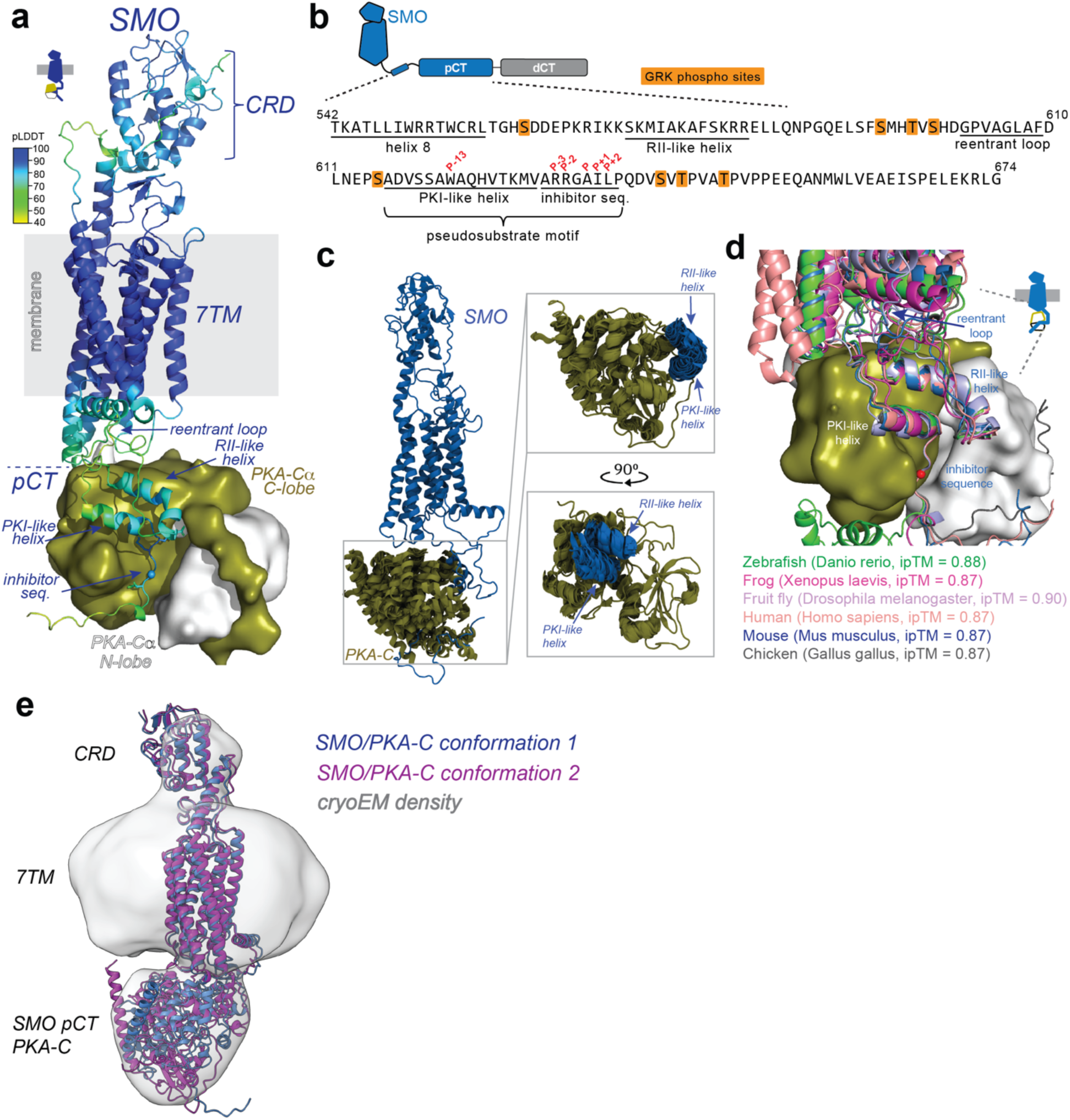
AlphaFold modeling of the SMO / PKA-C complex. **a,** AlphaFold 3 model of phosphorylated mouse SMO in complex with mouse PKA-Cɑ (rendered as a surface with small N-lobe in white and large C-lobe in olive). Positions of the SMO extracellular cysteine rich domain (CRD), transmembrane spanning domain (7TM), and proximal cytoplasmic tail (pCT) are indicated, and approximate location of membrane is shown in gray. The PKI-like helix, inhibitor sequence (seq), RII-like helix, and reentrant loop are labeled (see main text). The AlphaFold 3 predicted local confidence (pLDDT) score (0-100, with higher values indicating greater confidence) is represented by a color scale (upper left). The turquoise sphere in SMO’s inhibitor sequence represents the P-site. **b,** Sequence of SMO helix 8 and pCT. Secondary structural elements are underlined, GRK2/3 phosphorylation sites are highlighted in orange, and key amino acids in the pseudosubstrate motif are labeled in red. **c,** Structural snapshots of the complex between phosphorylated SMO (blue) and PKA-C (olive) as assessed by molecular dynamics (MD) simulations (one snapshot every 300 ns, 3 μs of simulation time). Each frame is aligned on the PKA-C subunit. Left: overall view of the complex, Right: zoomed-in view of the interface between PKA-C and SMO, highlighting the stability of the PKI-like helix, inhibitor sequence, and RII-like helix. **d,** AlphaFold3 models of SMO / PKA-C complexes from the indicated species, aligned on PKA-C from each complex. **e,** Docking of two AlphaFold 2.3.0 models (see Supplementary Discussion 2) into a low-resolution cryoEM map obtained from a BS3-crosslinked SMO / PKA-C sample in a GDN detergent micelle. Conformation 1 and conformation 2 are colored blue and dark magenta, respectively. CryoEM density is shown in gray and displayed at a contour level of 0.1.

Whereas some IDRs appear to be constitutively unstructured, others display conditional folding - they are unstructured in their *apo*, unmodified forms, but adopt partially or fully structured conformations upon post-translational modification and/or binding to other proteins^34–36^. Nevertheless, even the conditionally folded states of IDRs are challenging to characterize using empirical structural methods, due to their conformational heterogeneity^33–35,37^. Remarkably, AlphaFold, which derives structural models using multiple sequence alignments (MSAs) and coevolutionary analysis^44,45^, is well-suited to analysis of conditionally folded IDRs, as it often predicts them in their conditionally folded states, even when the requisite post-translational modifications and/or binding partners are not included in the model^46,47^ **(Supplementary Discussion 1)**. Intriguingly, prior AlphaFold analysis suggests that the SMO pCT may be a conditionally folding IDR, as the full-length SMO model available from the EBI database of AlphaFold structure predictions^44,45^ shows several helical regions in the pCT **(Extended Data Fig. 1b)**. We reasoned that AlphaFold’s potential to extract a physiological, conditionally folded SMO pCT state from a possibly vast and complex conformational ensemble may provide structural insights into the SMO / PKA-C interaction.

We used AlphaFold to model murine SMO bound to PKA-Cɑ (the best-studied and most ubiquitously expressed PKA-C isoform). We used AlphaFold 2.3.0 (also known as AlphaFold Multimer)^48^ to generate initial models but ultimately switched to AlphaFold 3 (which became available in the final stages of this project)^49^, since version 3 enabled us to explicitly model SMO phosphorylation sites as well as ATP and magnesium ions, all essential for SMO / PKA-C interactions^13–15^. The models produced by AlphaFold 2.3.0 and AlphaFold 3 are very similar, likely because the contributions from GRK2/3 phosphorylation are already reflected in the sequence-structure relationships in the MSAs^46^ **(Supplementary Discussion 1)**. AlphaFold 2.3.0 also predicts a second type of SMO / PKA-C model **(Extended Data Fig. 1c)**, which has a slightly lower interface predicted template modeling (ipTM, see below) score and may represent an alternative conformation of the complex **(Supplementary Discussion 2**; see also **Fig. 6, Extended Data Fig. 11a**). Hereafter we refer to the AlphaFold 3 model unless stated otherwise.

In the AlphaFold model, the phosphorylated SMO (pSMO) / PKA-C complex consists of an extended binding interface in which the SMO pCT and intracellular loops (ICLs) cradle PKA-Cs C-lobe and active-site cleft, with the kinase’s C-lobe pointing upward toward the SMO 7TM domain **(Fig. 1a)**. The complex harbors several key features **(Fig. 1a)**. Within the pCT, the pseudosubstrate motif consists of an inhibitor sequence that engages the PKA-C active site, immediately preceded by an amphipathic helix (615-630) which we refer to as the “PKI-like helix” **(Fig. 1a, b)** for its resemblance to one found in PKI proteins^26,50^. The PKI-like helix interacts at an approximately 30° angle with a second, longer amphipathic helix within the SMO pCT, which we designate the “RII-like helix” **(Fig. 1a, b)**, as it structurally mimics features of the PKA-RII holoenzyme^32^ (see below). The region between these helices contains several GRK2/3 phosphorylation sites along with a sequence, which we call the “reentrant loop” (602-609), that interacts intramolecularly with a cavity formed by transmembrane helices 3, 6, and 7 of the SMO 7TM domain **(Fig. 1a, b)**. Finally, other portions of the helices and intracellular loops from the SMO 7TM domain interact with PKA-C’s C-lobe, while the kinase’s N-terminal helical domain points towards the membrane **(Extended Data Fig. 1d)**.

Several observations indicate that our AlphaFold model accurately reflects the structure of the SMO / PKA-C complex, particularly in key interface-forming regions. First, the model agrees with prior empirical structures of PKA-Cɑ^31,32,50,51^ and the SMO extracellular and 7-transmembrane (7TM) domains^7,11,42,43^ **(Extended Data Fig. 1d)**. Second, the highest-ranked model scored 0.87 in AlphaFold’s ipTM measurements, indicating high global confidence^48^ in the position of the SMO / PKA-C interface. The model also scored >70 in predicted local distance difference test (pLDDT) and displayed low predicted aligned error (PAE) values within the PKI-like helix, inhibitor sequence, and RII-like helix, indicating high local confidence^52,53^ in the locations of residues within these key regions of the SMO / PKA-C interface **(Fig. 1a)**. Third, these same regions are stable throughout extended (3 µs) all-atom molecular dynamics simulations **(Fig. 1c)**, indicating that the AlphaFold model captures an energetically favorable mode of SMO / PKA-C interaction. Finally, the overall binding configuration of the SMO / PKA-C interface, as well as the underlying sequence and secondary structural elements, are conserved across metazoan evolution **(Fig. 1d, Extended Data Fig. 2a**).

To directly visualize the SMO / PKA-C complex, we undertook single-particle cryoEM studies of the complex purified and reconstituted in GDN detergent. Initial datasets revealed that the complex was unstable under standard conditions, so we employed bis(sulfosuccinimidyl)suberate (BS3), an amine-reactive crosslinker widely utilized in cryoEM studies^54–58^ to form intermolecular crosslinks and stabilize the complex **(Extended Data Fig. 3a**). Following 2D and 3D classification, we identified a promising class that produced a reconstruction at 8.59 Å nominal resolution **(Extended Data Fig. 3b–f)**. The resulting map revealed a large central ellipsoid corresponding to the GDN micelle, with two outwardly protruding lobes—one small and one large. Notably, our SMO / PKA-C AlphaFold models could be readily docked into this cryoEM density **(Fig. 1e)**. Although the resolution is insufficient to visualize main chain or side chain features, rigid-body fitting of the AlphaFold models demonstrated a good overall match to the experimental density. The low resolution of our reconstruction does not permit us to distinguish between the main vs. alternative SMO / PKA-C conformations predicted by AlphaFold. Nevertheless, the cryoEM map is consistent with the overall topology and relative positioning of SMO and PKA-C in our AlphaFold models. These empirical structural data demonstrate that our AlphaFold models are an accurate representation of the SMO / PKA-C interaction and a useful starting point for biochemical and functional characterization of this complex.

### A central role for the SMO pseudosubstrate motif in the SMO / PKA-C complex

In our model, the SMO pseudosubstrate motif forms the linchpin of the SMO / PKA-C interaction, with the inhibitor sequence adopting a canonical orientation sandwiched between the kinase’s N and C lobes^50^. Like other PKA-C inhibitors^29,31,32,50^, the SMO pseudosubstrate phosphorylation site (P-site) alanine, P-2 and P-3 arginines, and P+1 isoleucine residues interact extensively with the PKA-C active site cleft (**Fig. 2a**). The SMO inhibitor sequence is stabilized by hydrophobic packing of the P+1 isoleucine (SMO I636) against PKA-C L198, P202, and L205, and the P+2 leucine, P+3 proline, and P+6 valine (SMO L637, P638, and V641) against PKA-C L82 and F54 (**Fig. 2a**). In addition, SMO residues W622, V626, and M629, located at the N-terminus of the pseudosubstrate motif in the PKI-like helix, hydrophobically pack against Y235 and F239 within the PKA-C ɑF-ɑG region, resembling an analogous set of interactions that occur in PKI proteins^29,50^ (**Fig. 2a**). Consistent with our AlphaFold model, the P-2 and P-3 arginines, P-site alanine, and P-13 tryptophan (W622) are essential for SMO / PKA-C interactions and for Hh signal transduction based on mutational analysis in cultured cell and *in vivo* systems^14^.

**Figure 2:**
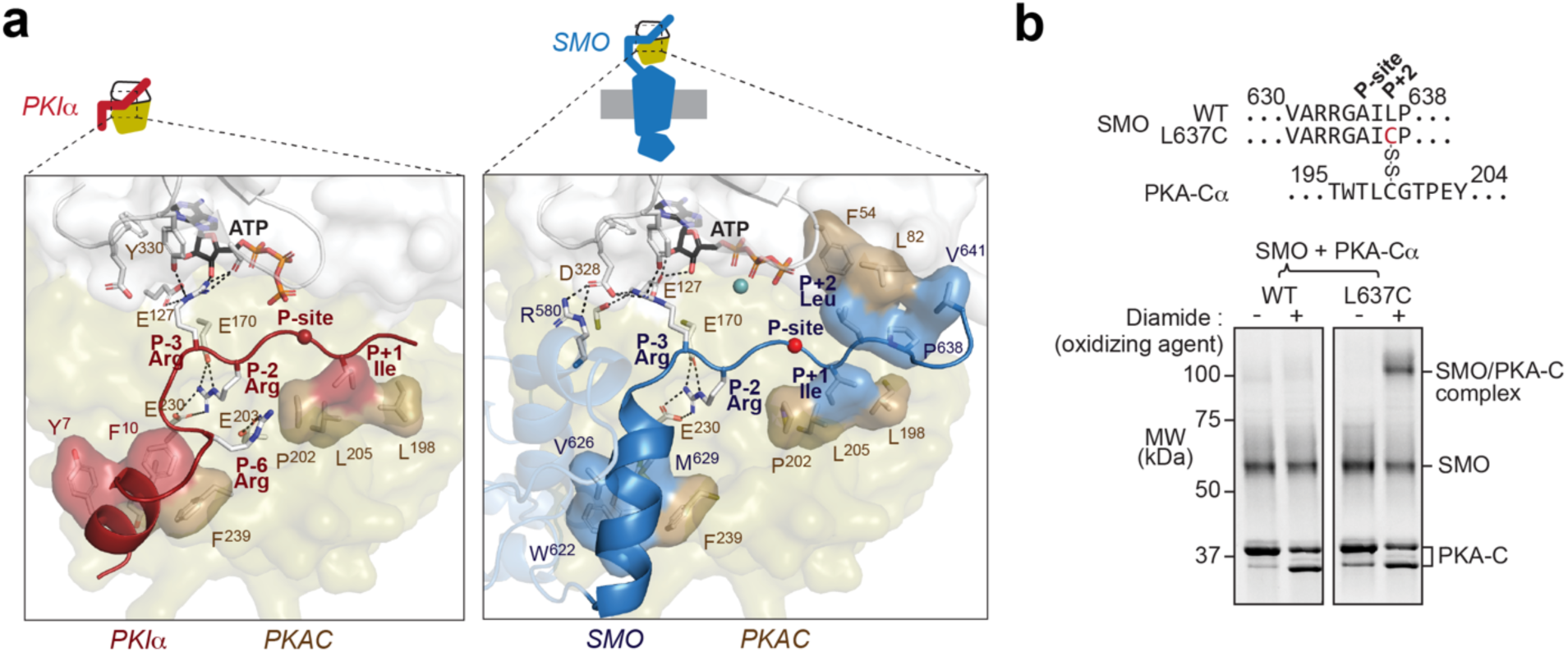
A central role for the SMO pseudosubstrate motif in the SMO / PKA-C complex. **a,** Left: empirical structure of PKIɑ(5-24) (PDB: 1ATP, red) in complex with PKA-C (olive/white as in Fig. 1a). Right: AlphaFold model of phosphorylated SMO (blue) in complex with PKA-C. In this and all subsequent figures, SMO cartoon (top) represents the orientation of the SMO / PKA-C complex (in this case with SMO at the bottom and PKA-C at the top, with the PKA-C N-lobe pointing upwards and the C-lobe pointing downwards). Locations of residues discussed in the text are indicated. Note salt bridge between the P-6 arginine in PKIɑ and PKA-C E203 (left); this interaction is absent in SMO (right). **b,** SDS-PAGE analysis of purified wild type (WT) or L637C mutant SMO incubated with PKA-C treated with or without the oxidizing agent diamide (to induce disulfide bond formation). Sequence alignments above the gel image show the mutation present in SMO and the location of the disulfide bond between SMO L637C and PKA-C C199. The endogenous cysteine in the SMO intracellular domain (C554) in wild-type SMO provides a negative control for nonspecific SMO / PKA-C disulfide bond formation. Location of SMO, PKA-C, and the disulfide-trapped SMO / PKA-C complex are indicated at right. PKA-C runs as a doublet following diamide treatment, due to formation of an intramolecular disulfide bond between C199 and C343 as shown previously^62^.

To further evaluate the SMO / PKA-C binding mode in our models, we conducted disulfide trapping studies. In PKA-RII holoenzymes, a cysteine at the P+2 position in the PKA-R subunit forms a disulfide bond with PKA-C C199 under oxidizing conditions^59,60^. If the SMO and PKA-RII inhibitor sequences bind similarly to PKA-C, then substitution of the SMO P+2 leucine with cysteine (L637C) should readily produce a disulfide bond between SMO and PKA-C C199, similar to the naturally occurring P+2 cysteine in PKA-RII. This was indeed observed (**Fig. 2b, Extended Data Fig. 2b, c**), providing strong support for this region of the SMO / PKA-C binding configuration captured in our AlphaFold model.

Whereas PKI proteins operate as constitutive PKA-C inhibitors^26,28^, only the active, GRK2/3-phosphorylated conformation of SMO can bind and inhibit PKA-C under cellular conditions^13,15^. Accordingly, peptides encompassing the minimal active site-binding portion of PKIɑ bind PKA-C with low nanomolar affinity^61^, but a peptide encompassing the corresponding region of SMO binds nearly three orders of magnitude more weakly^14^. The PKI-like helix in SMO is elongated and reorientated in our models compared to the one in PKIɑ, resulting in loss of a salt bridge essential for high-affinity binding (involving the PKIɑ P-6 arginine and PKA-C E203 in the PKIɑ / PKA-C complex)^50^ (**Fig. 2a**). This reorientation likely explains why the pseudosubstrate motif in SMO binds PKA-C less efficiently than does its counterpart in PKIɑ.

Thus our AlphaFold model provides a high-confidence structural prediction of the SMO / PKA-C complex structure and establishes the central role of SMO’s PKA pseudosubstrate motif in mediating this interaction.

### SMO structurally mimics the PKA-RIIβ holoenzyme

To bind PKA-C with high affinity, PKA-R subunits combine a “central” interaction at the PKA-C active-site cleft with “auxiliary” interactions engaging additional PKA-C surfaces^2,25,27^. In the tetrameric PKA-RIIβ holoenzyme (RIIβ_2_-C_2_), the inhibitor sequence from one PKA-RIIβ subunit binds the PKA-C active-site cleft, while the β4-β5 loop from the other PKA-RIIβ subunit engages the PKA-C hinge region (between the kinase’s N- and C-lobes)^63^ and “FDDY” motif which is essential for ATP binding^32^ (**Fig. 3a**). Together, these central and auxiliary interactions lock PKA-C into a closed conformation and interrupt catalysis^2,25,27^.

**Figure 3:**
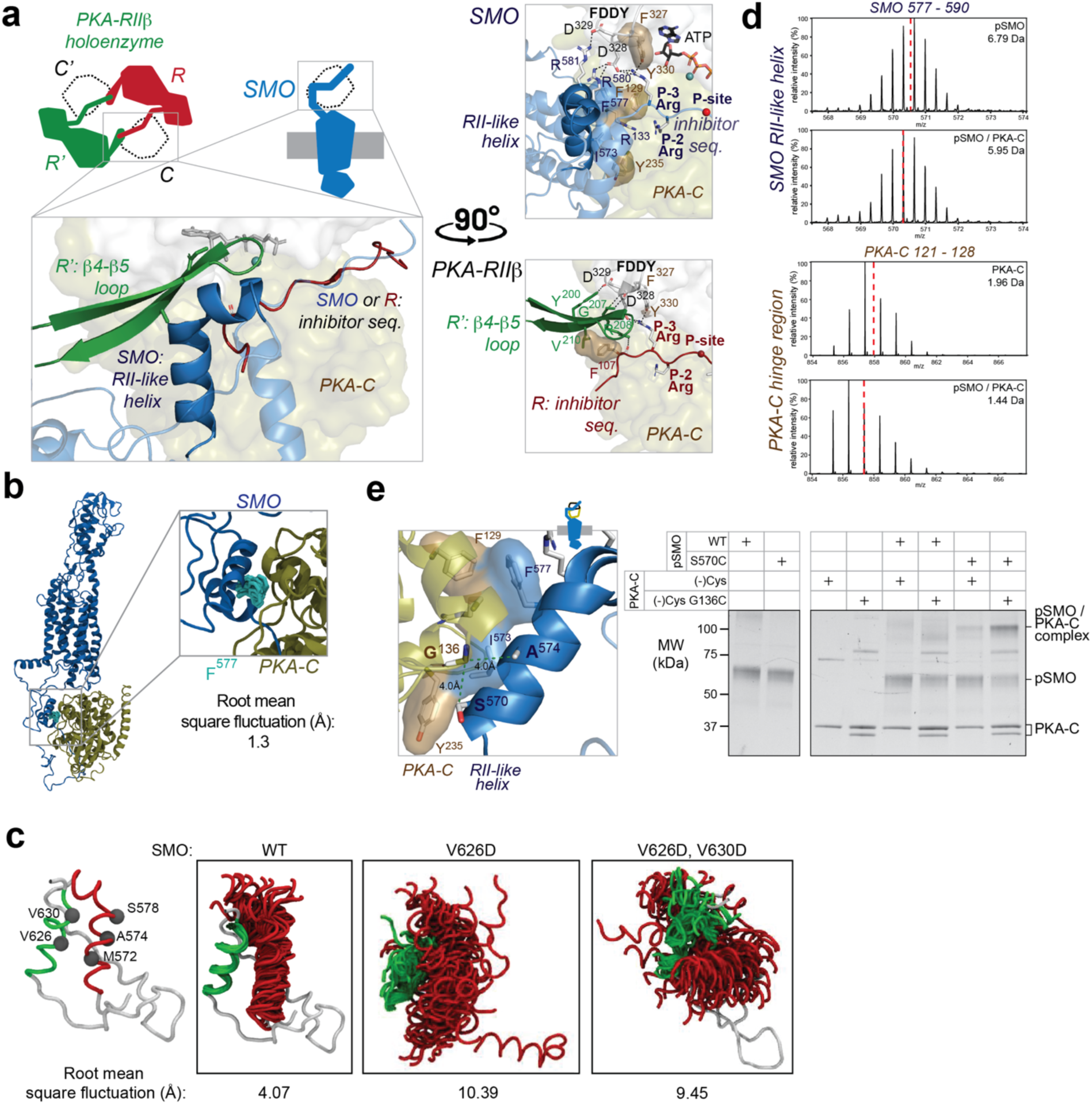
Structural mimicry of the PKA-RIIβ holoenzyme by a SMO amphipathic helix. **a,** Top: cartoon of tetrameric PKA-RIIβ holoenzyme (RIIβ_2_-C_2_, left cartoon) in which each PKA-C subunit (C, dotted outline) contacts the inhibitor sequence from one PKA-R subunit (R, red) and the β4/β5 loop from the other PKA-R subunit (R’, green), and vice versa. Bottom: overlay between PKA-RIIβ holoenzyme empirical structure (PDB: 3TNP) and phosphorylated SMO / PKA-C AlphaFold 3 model aligned on the PKA-C in each complex. SMO is shown in blue, with the RII-like helix in the foreground (dark blue) and additional SMO sequences, including the inhibitor sequence and PKI-like helix, in the background (light blue). The SMO and PKA-RIIβ complexes are shown separately at right, with key residues and motifs indicated as described in the main text. **b,** MD structural snapshots of SMO F577 (light blue, one snapshot every 150 ns, 3 μs of simulation time) within the SMO C-tail (dark blue), when aligning the simulation frames with PKA-C (brown). **c,** MD structural snapshots (one snapshot every 30 ns, 1.5 μs of simulation time) show the positions of the PKI-like helix (615-630, green) and RII-like helix (570-581, red) during simulations of wild-type SMO (WT) or the indicated SMO mutants. Overall root-mean square fluctuations (in Å) for the residues indicated in the model at left are shown below each simulation. **d,** Representative HDX-MS mass spectral envelope (deuterium exchange time t_ex_=10 min) of peptides in the indicated region in phosphorylated SMO (pSMO) or PKA-C alone (top spectrum in each pair), or in complex with PKA-C (bottom spectrum in each pair). The deuterium uptake is shown for each spectrum on the top right. Centroids are indicated by red dashed lines. SMO peptide = 577-590, and PKA-C peptide = 121-128. **e,** SDS-PAGE analysis of diamide-induced disulfide trapping between pSMO S570C and PKA-C G136C in a minimal-cysteine ((-)Cys) background (see Methods). Wild-type (WT) SMO (harboring an endogenous cysteine in helix 8, see Fig. 1) or (-)Cys PKA-C, serve as negative controls. Locations of SMO S570, A574, and PKA-C G136 are indicated in the cartoon at left. See Fig. 5e for quantification.

Surprisingly, a 12 amino acid SMO amphipathic helix (residues 570-581 in mouse SMO), which we term the “RII-like helix”, mimics key structural features of both PKA-R subunits in the PKA-RIIβ holoenzyme. Although this helix displays no obvious sequence or structural homology to PKA-RIIβ, our model reveals that it binds to the same regions engaged by the β4-β5 loop of one PKA-RIIβ subunit and the backside of the inhibitor sequence from the other^32^ (**Fig. 3a**). Furthermore, an analogous pattern of hydrophobic and hydrogen bonding interactions underlies both SMO / PKA-C and PKA-R / PKA-C complexes. Specifically, SMO F577 hydrophobically packs against PKA-C F129 and R133, mirroring the hydrophobic interactions of F107 at the P-5 site from the PKA-RIIβ inhibitor sequence and V210 from the β4-β5 loop^32^ (**Fig. 3a**). SMO R580 and R581 also form hydrogen bonds with PKA-C D328 and D329 from the FDDY motif, mimicking interactions between PKA-RIIβ Y200, G207, and R208 with these same PKA-C residues^32^ (**Fig. 3a**). A cation-pi interaction between SMO R580 and SMO F577 is expected to enhance these ionic interactions (**Fig. 3a**). Additional residues contributing to this region of the SMO / PKA-C complex are summarized in Fig. 3a. Thus, despite distinct primary sequences, secondary structural elements, and binding stoichiometries, SMO and PKA-RIIβ utilize remarkably similar central and auxiliary interactions to bind and inhibit PKA-C.

The interfaces between the SMO PKI-like helix, inhibitor sequence, RII-like helix, and the PKA-C hinge region and active site cleft remain stable during all-atom MD simulations of the SMO / PKA-C complex, with SMO F577 exhibiting exceptional stability (**Fig. 3b, c**). These observations are indicative of an energetically favorable binding configuration. To empirically determine which parts of SMO and PKA-C participate in complex formation, we employed amide hydrogen/deuterium exchange mass spectrometry (HDX-MS, **Extended Data Fig. 4, 5a-c**)^64–67^. HDX-MS monitors changes in hydrogen bonding and solvent accessibility of backbone amides in SMO and PKA-C upon complex formation, thereby providing a peptide-level readout of conformational changes in each protein upon complex formation. Initial attempts to purify phosphorylated SMO / PKA-C complexes for HDX-MS were hampered by instability of the complex during gel filtration chromatography (data not shown). Inspired by strategies to stabilize GPCR signaling complexes for structural studies^68–70^, we disulfide-trapped the SMO / PKA-C complex at the previously validated interface between the SMO pseudosubstrate motif and PKA-C active site cleft (SMO L637C + PKA-C C199, **Fig. 2b)**; this highly specific disulfide bond is expected to stabilize the complex’s central interaction while allowing other peripheral interactions to form freely. AlphaFold modeling indicated that this disulfide trap is not likely to extensively alter peripheral regions of the SMO / PKA-C complex **(Extended Data Fig. 5d)**. HDX-MS analysis of the disulfide-trapped, phosphorylated SMO / PKA-C complex revealed deuterium exchange protection in the SMO pseudosubstrate motif and the PKA-C active site cleft relative to the uncomplexed proteins **(Extended Data Fig. 5e, f; Supplementary Discussion 3).** The SMO RII-like helix (577-590), as well as the PKA-C hinge region (122-128), also showed significant deuterium exchange protection (>0.5 Da) in the complex (**Fig. 3d, Extended Data Fig. 6a**). Overall, our empirical findings are consistent with the SMO RII-like helix binding the PKA-C C-lobe as captured in our model of the complex.

To directly evaluate the binding between the SMO RII-like helix and the PKA-C hinge region with single-residue precision, we conducted additional disulfide trapping studies. In our model, SMO S570 and A574 on the hydrophobic face of the RII-like helix reside within 2.7 and 4.4 Å of PKA-C G136, respectively. Accordingly, purified, phosphorylated SMO S570C or A574C formed a specific disulfide bond with PKA-C G136C (**Fig. 3e, Extended Data Fig. 6b**), demonstrating that the SMO RII-like helix directly contacts the PKA-C hinge region, consistent with the AlphaFold model.

To assess the impact of the SMO RII-like helix on SMO / PKA-C interactions and Hh signal transduction, we utilized mutational analysis. A previous systematic scanning mutagenesis study of SMO intracellular regions identified eight amino acids that, when mutated individually to alanine (or to glycine, for alanines in the wild-type sequence), prevent Hh-induced activation of a GLI transcriptional reporter, via an unknown mechanism^71^. Notably, six out of eight of these residues are within the SMO RII-like helix, and four of the six (S570, I573, F577, R580) directly engage PKA-C in our AlphaFold model (**Fig. 4a**). We confirmed these observations by showing that alanine substitution of I573 and F577, either individually (I573A or F577A) or in combination with R580A (IFR → AAA), abolished GLI reporter activation by native Sonic hedgehog proteins (ShhN) or the direct SMO agonist SAG21k (**Fig. 4a, Extended Data Fig. 6c**). The remaining two essential residues in the RII-like helix (A574, A576)^71^ do not face PKA-C, but the mutations analyzed previously (A574G, A576G)^71^ likely hindered formation of the RII-like helix, as glycine is a known helix-destabilizing residue. To further evaluate the requirement for helicity, we engineered a helix-breaking mutation at SMO K575, a non-PKA-C-interacting residue within the RII-like helix. Whereas alanine substitution of this residue only modestly (<50%) decreased GLI reporter activation^71^, substitution with proline, a potent helix-disrupting residue, abolished GLI transcriptional activation (**Fig. 4b, Extended Data Fig. 6c**). Finally, the RII-like helix is stabilized via a hydrophobic interface with the PKI-like helix **(Extended Data Fig. 6d**), and substitution of one or both valines underlying this interface with a hydrophilic residue, aspartate (SMO V626D and V630D), destabilized the interface in MD simulations (**Fig. 3c**) and eliminated GLI transcriptional activation in cells (**Fig. 4b, Extended Data Fig. 6c**). Control experiments established that the inability of these mutants to signal cannot be explained by defects in SMO expression, ciliary localization, or agonist-induced GRK2/3 phosphorylation **(Extended Data Fig. 6e, f**), consistent with previous findings^13,71,72^.

**Figure 4:**
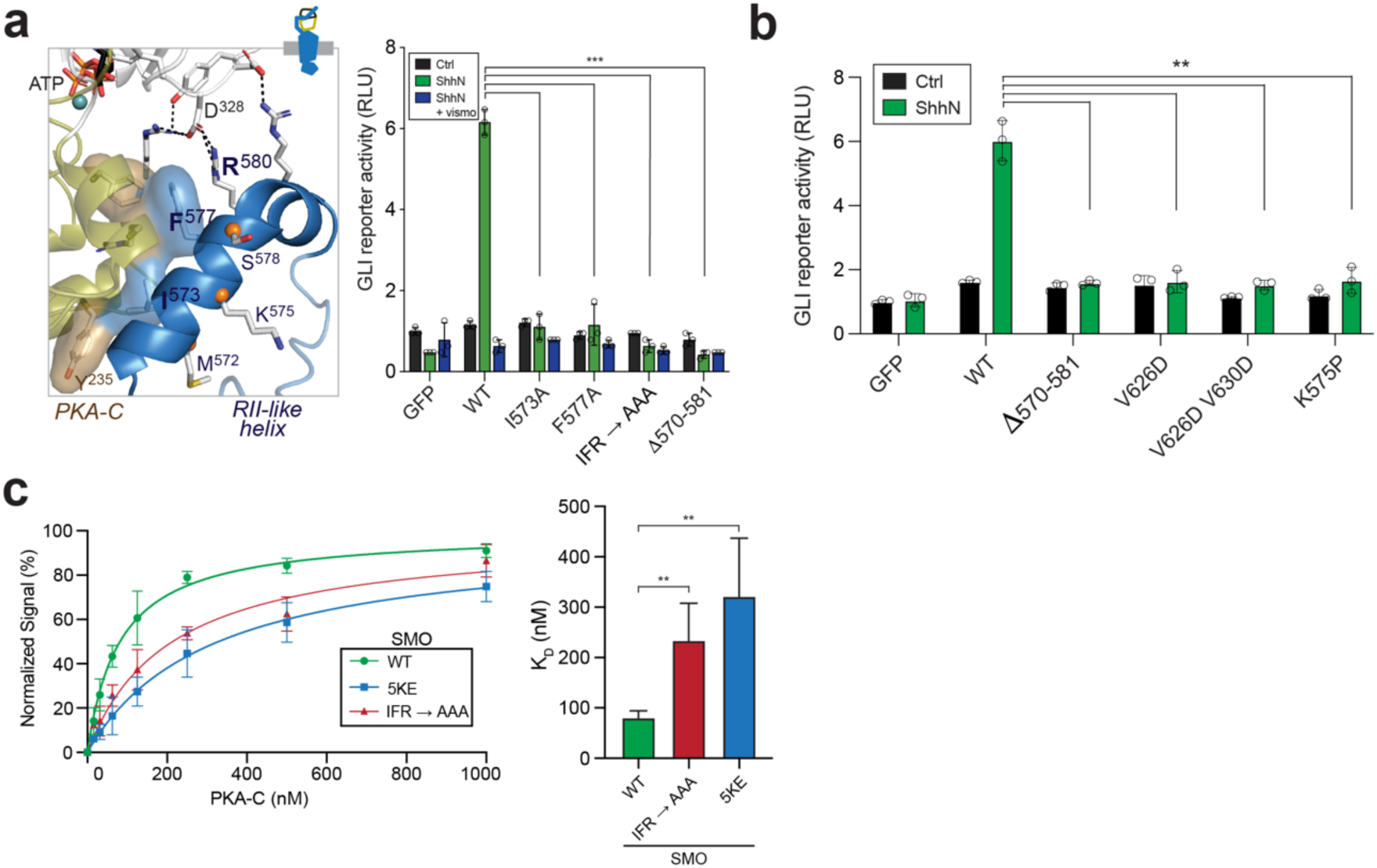
The SMO RII-like helix is essential for Hh signal transduction. **a,** GLI transcriptional reporter assay in *Smo^-/-^* murine embryonic fibroblasts (MEFs) transfected with GFP (negative control), WT SMO, or the indicated SMO mutant constructs. Cells were treated with conditioned medium containing the N-terminal signaling domain of Sonic hedgehog (ShhN) alone (green), or in the presence of the SMO inverse agonist vismodegib (ShhN + vismo, blue), or control conditioned medium lacking ShhN (black). Data represents the mean +/- s.d., n=3 independent biological samples. Locations of SMO residues are indicated in the cartoon at left. **b,** A second panel of SMO mutants were analyzed as in (**a**). Data represents the mean +/- s.d., n=3 independent biological samples. **c,** Left: Steady-state SPR analysis of binding interactions between PKA-C and wild-type SMO or the indicated SMO mutants reconstituted into nanodiscs (see Figure 5 for definition and functional characterization of the 5KE mutant); Right: K_D_ determinations for the experiments shown at left (n=4 measurements per condition).

A parsimonious explanation for the Hh signaling loss-of-function mutations in the SMO RII-like helix and PKI-like helix is that these mutations all block Hh signal transduction by compromising a key binding interface formed by the SMO RII-like helix and PKA-C hinge region. Consistent with this hypothesis, the region of SMO encompassing the RII-like helix (570-581) is critical for interactions with PKA-C in cultured cell bioluminescence resonance energy transfer (BRET) assays^13^. To quantitatively assess the role of the SMO RII-like helix in mediating SMO / PKA-C interactions, we developed a surface plasmon resonance (SPR) assay to measure the affinity of PKA-C for near-full-length, GRK2/3-phosphorylated SMO (pSMO) reconstituted into nanodiscs *in vitro*. In this assay, wild-type pSMO bound PKA-C with high affinity (K_D_ = 82 +/- 11 nM) (**Fig. 4c, Extended Data Fig. 7; Supplementary Discussion 4)**, while the IFR→ AAA mutation significantly weakened the interaction (K_D_ 222 +/- 63 nM) (**Fig. 4c, Extended Data Fig. 7**). Although the effects of the mutation were less pronounced in the *in vitro* binding assay compared to the cell-based GLI transcriptional reporter assay (**Fig. 4a**), possibly arising from differences in the dynamic range of each assay or in the level of SMO phosphorylation in each experiment **(Supplementary Discussion 4)**, these results are consistent with the role of the RII-like helix in mediating the SMO / PKA-C interaction.

In sum, our AlphaFold modeling, HDX-MS, disulfide trapping, and mutational analyses demonstrate that SMO binds PKA-C by structurally mimicking the PKA-RIIβ subunit in PKA holoenzymes.

### Phosphorylation generates new structural elements that stabilize the SMO / PKA-C complex

GRK2/3 dramatically enhances the SMO / PKA-C interaction^15^ by phosphorylating active SMO at eight conserved serines and threonines within the pCT that we mapped via mass spectrometry^13,15^ **(Extended Data Fig. 8a**). Five of these residues (pS560, pS594, pT597, pS599, pS615) reside in the N-terminal portion of the pCT, preceding the PKI-like helix, while three (pS642, pT644, pT648) are located C-terminal to the inhibitor sequence (**Fig. 1b**). Our AlphaFold models immediately suggest mechanisms by which phosphorylation in each of these regions enhances the SMO / PKA-C interaction.

The five N-terminal phosphorylated residues localize near a stretch of seven conserved lysines and arginines (K/R) (**Fig. 5a, Extended Data Fig. 8b**). We hypothesize that phosphorylation fosters electrostatic interactions between the negatively charged phospho-serines / threonines (pS/pT) and positively charged K/Rs. This will constrain the otherwise disordered SMO cytoplasmic tail into a more compact, ordered state conducive to forming key secondary structures in the PKA-C binding interface, namely the RII-like helix and the PKI-like helix. Thus, we postulate that a phosphorylation-driven intramolecular electrostatic switch in SMO drives SMO / PKA-C interactions.

**Figure 5:**
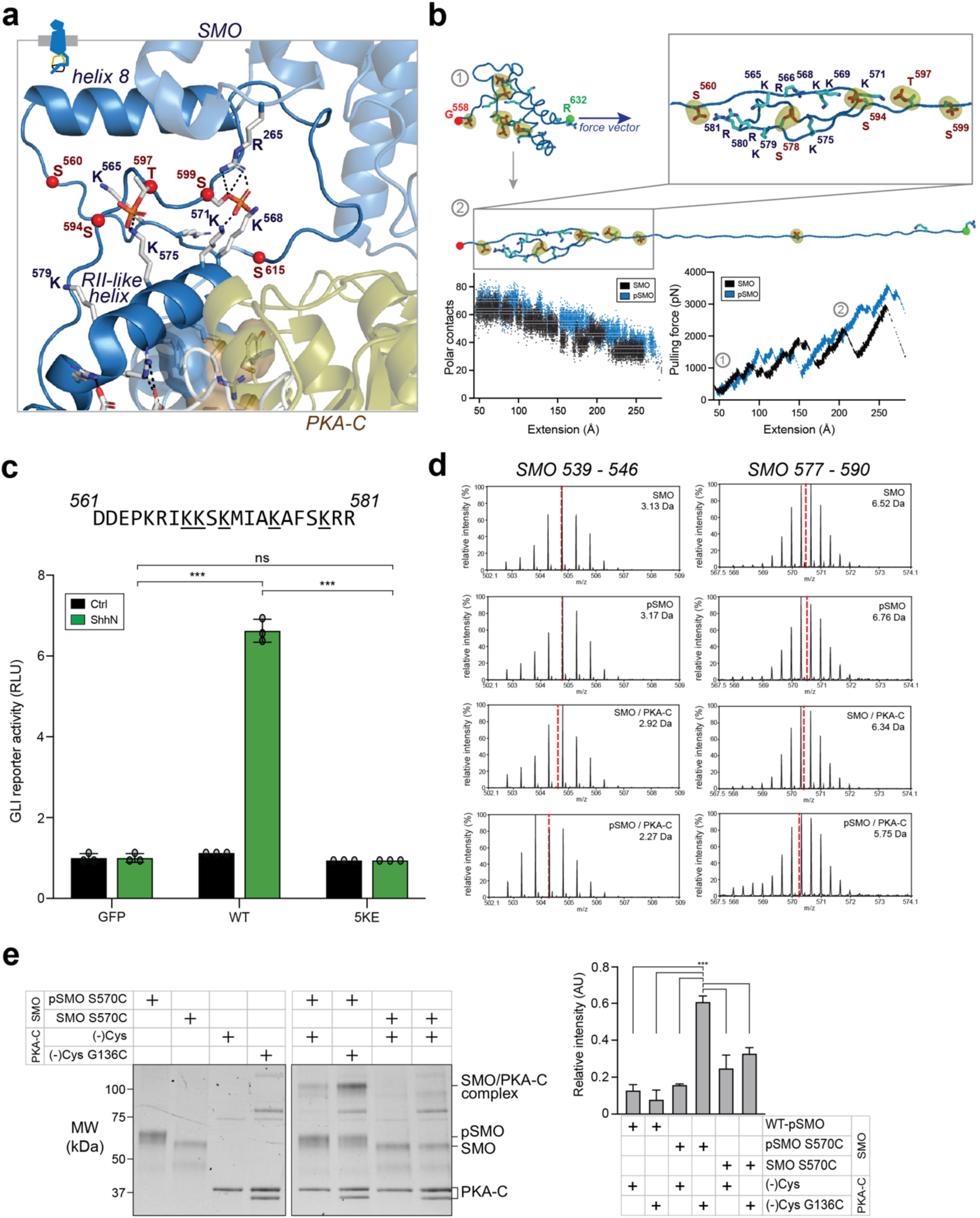
SMO phosphorylation generates new structural elements that stabilize the PKA-C complex. **a,** AlphaFold model of the SMO / PKA-C complex (SMO in blue, PKA-C in olive/white), with GRK2/3-phosphorylated S/T residues (red) and conserved K/R residues highlighted in the SMO pCT. SMO helix 8 and the RII-like helix are indicated for orientation. **b,** Stability of the SMO pCT in its phosphorylated vs nonphosphoryated states was assessed in MD simulations by applying a force vector (indicated by the direction of the blue arrow) to unfold the pCT, then quantifying the force required to achieve a constant velocity of unfolding^74,75^, monitored as the distance between the red and green spheres. Top: Beginning (1) and ending (2) states for the MD simulation (states are marked on the bottom right graph) with a vector indicating the direction of applied force. Bottom: Graphs showing “extension” (distance between the red and green spheres at the ends of SMO) on the X-axes and either “polar contacts” or “pulling force” on the Y-axis for phosphorylated (blue) vs nonphosphorylated (black) SMO, showing that SMO phosphorylation increases the number of intramolecular polar contacts leading to an increased force required to unfold the SMO pCT. **c,** Wild-type SMO or the 5KE mutant (underlined K/R residues mutated to E) were analyzed via a GLI transcriptional reporter assay in *Smo^-/-^* MEFs, performed as in Fig. 4a, b. **d,** Representative HDX-MS mass spectral envelope (t_ex_= 5 min) of the indicated peptide in SMO helix 8 (539-546) or the RII-like helix (577-590) in the nonphosphorylated vs phosphorylated SMO / PKA-C complex (SMO / PKA-C (3^rd^ row) vs pSMO / PKA-C (4^th^ row), respectively). The deuterium uptake is shown for each spectrum on the top right. Centroids are indicated by red dashed lines. **e,** Disulfide trapping of PKA-C wild-type or G136C (in a minimal cysteine ((-)Cys construct) with phosphorylated vs nonphosphorylated, SAG21k-bound SMO (pSMO vs SMO, respectively), as in Fig. 3e. Band intensity is quantified in the graph at right, and represents mean +/- standard deviation from 3 independent trials. Representative gel image corresponding to disulfide trapping of phosphorylated wild-type vs S570C pSMO is shown in Fig. 3e.

Several computational and experimental results support our proposed mechanism. First, using MD simulations to determine the stability of the SMO pCT in the presence or absence of phosphorylation, we found that interactions between the pS/pT and K/R residues create polar contacts that stabilize the conditionally folded state of the pCT (**Fig. 5b**). Second, mutation of K/R residues within this conserved SMO sequence stretch (5KE) reduced the affinity of SMO for PKA-C *in vitro* (**Fig. 4c, Extended Data Fig. 7**) and eliminated GLI reporter activation in cultured cells (**Fig. 5c, Extended Data Fig. 8c**), without blocking SMO expression, trafficking, or agonist-induced phosphorylation **(Extended Data Fig. 6e, f**). Thus, the K/R stretch in the SMO pCT is essential for SMO / PKA-C interactions and Hh signal transduction, paralleling findings for the GRK2/3 phosphorylation sites^13^ and PKA-C-interacting residues in the RII-like helix (**Fig. 4a, b**). Third, HDX-MS studies of phosphorylated vs. nonphosphorylated SMO / PKA-C complexes **(Supplementary Discussion 3)** revealed that phosphorylation leads to a pronounced deuterium exchange protection (>0.5 Da) in a peptide comprising the majority of the SMO RII-like helix (577-590), and another comprising most of the amphipathic helix 8 from the 7TM domain (539-546) (**Fig. 5d, Extended Data Fig. 6a, 9a-c)**, which runs parallel to the membrane and lies just above the pS/pT and K/R residues (**Fig. 5a**). Despite the high relative concentrations of SMO and PKA-C in these disulfide-trapped complexes, little to no protection occurs at SMO helix 8 or the RII-like helix unless SMO is phosphorylated, highlighting the profound importance of phosphorylation for stabilizing the SMO pCT and facilitating binding to PKA-C. Lastly, disulfide trapping between SMO S570C or A574C in the SMO RII-like helix and G136C in the PKA-C hinge region is substantially weaker when SMO is not GRK2/3-phosphorylated (**Fig. 5e, Extended Data Fig. 9d**), directly demonstrating that phosphorylation stabilizes an essential SMO / PKA-C binding interface.

Besides these five GRK2/3-phosphorylated residues, SMO also harbors three GRK2/3-phosphorylated residues C-terminal to the PKA-C pseudosubstrate motif. Similarly, the ryanodine receptor (RyR), a PKA-C substrate, has phosphorylation sites for Ca^2+^/calmodulin-dependent protein kinase 2 (CaMKII) located C-terminal to its PKA-C substrate motif **(Extended Data Fig. 9e**), and CaMKII phosphorylation enhances RyR / PKA-C interactions by promoting intermolecular hydrogen bonds and stabilizing PKA-C-interacting structural elements^73^. These findings suggest that GRK2/3 phosphorylation may utilize a similar strategy to enhance SMO / PKA-C interactions in this region. Intriguingly, this region of SMO is strongly protected upon GRK2/3 phosphorylation in HDX-MS studies, consistent with a disorder-to-order transition, and the protection is enhanced when PKA-C is present **(Extended Data Fig. 9e, f**). While our AlphaFold models are of limited confidence in this region of SMO, they hint that these GRK2/3-phosphorylated residues may enhance SMO’s ability to wrap around PKA-C **(Extended Data Fig. 9g**).

Taken together, our results demonstrate that SMO phosphorylation enables SMO / PKA-C binding by promoting intra- and intermolecular electrostatic interactions that stabilize conditionally folded regions within the SMO cytoplasmic domain, thereby remodeling it into a PKA-C-inhibiting conformation.

### Structural mimicry of canonical GPCR-effector interactions by the SMO / PKA-C complex

Thus far, our analysis of SMO / PKA-C interactions has focused on structural similarities between SMO and classical PKA-C inhibitors, namely PKI proteins and PKA-R subunits. However, the SMO / PKA-C complex engages in a second, parallel form of structural mimicry of canonical GPCRs in complex with conventional downstream effectors such as G proteins and β-arrestins^16,17,24^. Comparing our SMO / PKA-C model to an empirical structure of a prototypical GPCR bound to a β-arrestin^76^ (M2 muscarinic receptor (M2AchR) / β-arrestin1) reveals an overall architectural similarity between these complexes (**Fig. 6a**). Moreover, the SMO 7TM domain, as well as membrane lipids, contribute to the SMO / PKA-C interaction in a number of ways that echo GPCR-effector complexes.

**Figure 6:**
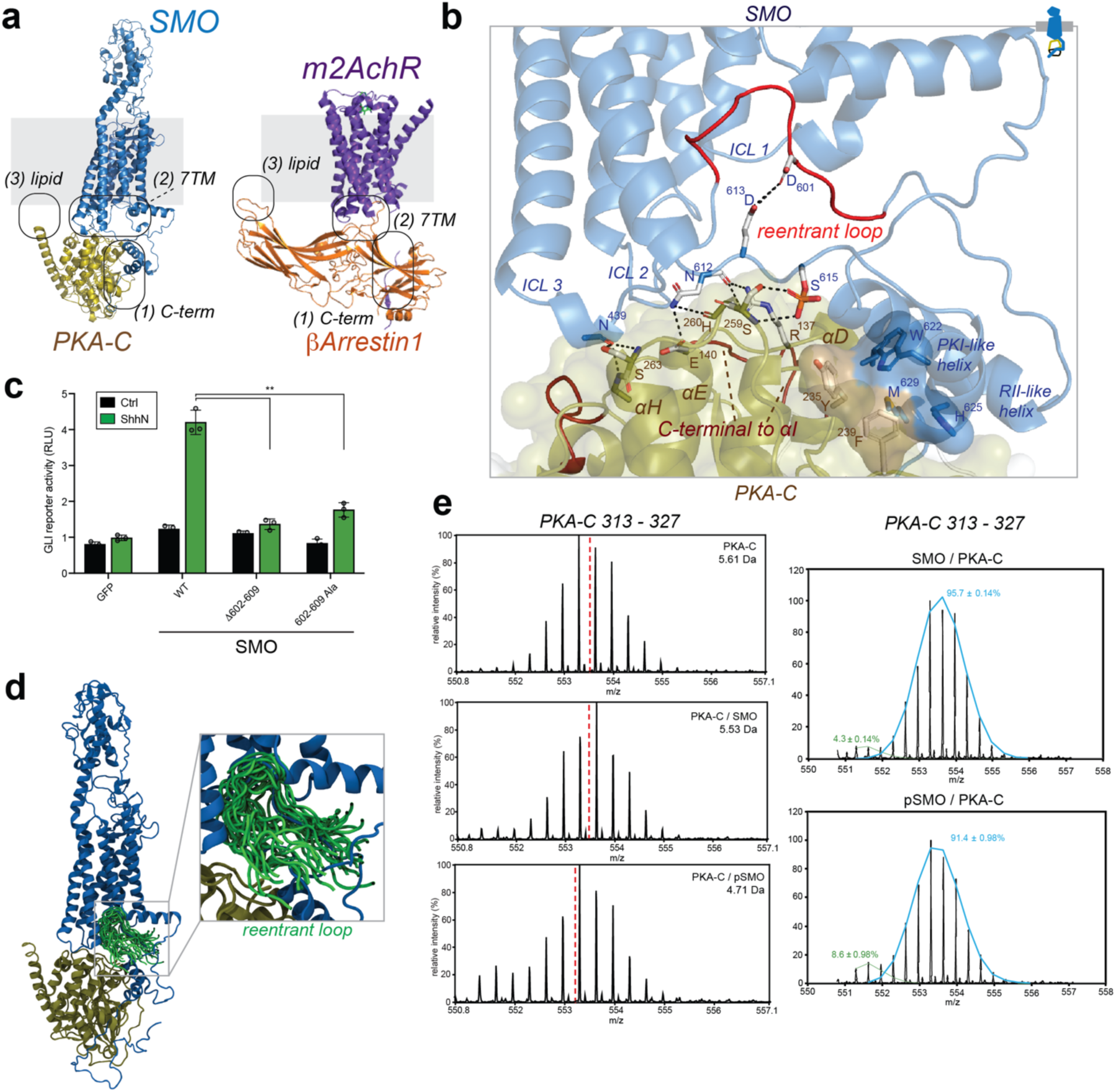
Structural mimicry of canonical GPCR-effector signaling complexes. **a,** Comparison of the SMO / PKA-C model (left) with the m2AchR / β-arrestin1 structure (right). Locations of interaction sites in the C-terminus (including the reentrant loop), 7TM domain (including ICLs), and membrane are indicated in each complex. **b,** The SMO 7TM interface. Residues in the αD-αE and αG-αH loops of PKA-C interacting with SMO ICL domains are indicated. The reentrant loop region of the SMO pCT (residues 602-609) is shown in red. **c,** GLI reporter assay for the indicated deletions or alanine substitutions in the SMO reentrant loop (see (**b**)). **d,** Structural snapshots of the complex between phosphorylated SMO (blue) and PKA-C (olive) (one snapshot every 300 ns, 3 μs of simulation time). Each frame is aligned on the PKA-C subunit. Reentrant loop region of SMO pCT is highlighted in green. **e,** HDX-MS data for region of PKA-C C-terminal to the αI helix (residues 313-327) in its apo (top), SMO-bound (middle), or pSMO-bound (bottom) forms, determined at t_ex_ = 10 min and plotted as in Fig. 3d. Bimodal deconvolution of the PKA-C 313-327 spectra revealed high-exchanging (blue) and low-exchanging (green) populations, suggesting two non-interconverting states of the SMO / PKA-C complex.

First, a region of the SMO pCT, termed the reentrant loop (602-609), inserts into a cavity at the intracellular face of the 7TM domain (**Fig. 6b**). This cavity is a nexus for protein-protein interactions in GPCRs – it opens selectively upon conformational activation of the GPCR 7TM domain, enabling helical elements or loops from G proteins^77,78^, GRKs^57,79^, β-arrestins^70,76,80^, and conformation-specific nanobodies^11,68,81^ to selectively engage the GPCR in its activated state **(Extended Data Fig. 10a**). In the β2-adrenergic receptor, a conventional GPCR, the receptor’s cytoplasmic tail interacts with its 7TM domain, which restricts access of G proteins to the 7TM cavity to dampen downstream signaling^39^. For SMO, in contrast, this reentrant loop / cavity interaction is expected to promote signaling to PKA-C, as it draws the pseudosubstrate motif and RII-like helix close together (**Fig. 6b**), into an optimal configuration to bind to the PKA-C hinge region and active-site cleft. Accordingly, deletion or alanine substitution of residues in the reentrant loop strongly inhibited GLI reporter activation (**Fig. 6c**), without substantially affecting expression, ciliary trafficking, or ability to undergo GRK2/3-mediated phosphorylation **(Extended Data Fig. 6e, f)**. Although AlphaFold’s pLDDT scores in this region (50-70) are modest, the interaction between the SMO reentrant loop and 7TM cavity remains throughout all-atom MD simulations (**Fig. 6d**), occurs in SMO / PKA-C complexes from multiple orthologs (**Fig. 1d**), and persists even when the SMO 7TM domain and reentrant loop are modeled as separate polypeptide sequences **(Extended Data Fig. 10b)**. These results underscore how the SMO 7TM cavity, like those in other GPCRs, enables critical interactions with downstream effectors.

Second, intracellular loops (ICLs) 1 and 2 of the SMO 7TM domain engage the surface of PKA-C’s C-lobe, similar to ICL interactions in canonical GPCR complexes with their effectors^70,76,77,79^ **(Extended Data Fig. 10c)**. In GPCR / β-arrestin complexes, the ICLs can bind to several distinct β-arrestin surfaces **(Extended Data Fig. 10c)**, leading to a range of conformations which may contribute to diverse signaling outcomes^16,17,82,83^. The SMO / PKA-C complex may exhibit similar conformational heterogeneity, as AlphaFold produces two classes of conformations of the SMO / PKA-C complex with the ICLs engaging distinct C-lobe interfaces **(Extended Data Fig. 10c)** involving the PKA-C αD-αE and αG-αH loops (**Fig. 6b, Extended Data Fig. 11a; Supplementary Discussion 2)**. These binding interfaces resemble docking sites those used by other PKA-C interactors, such as PKA-R’s cyclic nucleotide binding domains (CNBD)^31,51^ and the cystic fibrosis transmembrane regulator (CFTR) ion channel^84^; furthermore, the αG-αH loop functions as a hub for protein-protein interactions throughout the AGC kinase family^85^. HDX-MS studies revealed strong protection of the PKA-C αG-αH region in the SMO / PKA-C complex compared to PKA-C alone **(Extended Data Fig. 11a)**, supporting the formation of this interface in the SMO / PKA-C complex.

Third, our AlphaFold models suggest roles for lipids in SMO / PKA-C interactions. Lipids stabilize conventional GPCR signaling complexes at the membrane, in part by increasing local concentrations of the GPCR effector at the membrane inner leaflet ^76,79,86–91^. Membrane lipids may play analogous stabilizing roles in the SMO / PKA-C complex. In support of this hypothesis, HDX-MS studies revealed that a region of the PKA-C C-lobe (313-327) modeled by AlphaFold to face upward toward the membrane undergoes protection in the SMO / PKA-C complex (**Fig. 6e, Extended Data Fig. 11b**), suggestive of direct interactions with the membrane (or the detergent micelle in our *in vitro* systems). These interactions may be electrostatic in nature, as the SMO / PKA-C complex presents a highly electropositive surface near the membrane, favoring interactions with negatively charged phospholipid head groups **(Extended Data Fig. 11c**). Alternatively, or in addition, PKA-C is N-terminally myristylated, and this moiety can be mobilized from its hydrophobic binding pocket between the αA helix and the C lobe to promote membrane interactions^84,92–97^. Although AlphaFold cannot yet incorporate membranes or N-myristyl groups, it is intriguing that the PKA-C N-terminus is oriented toward the membrane in our AlphaFold models, which may facilitate membrane insertion of the myristyl moiety (**Fig. 6a**). Thus, lipid interactions may stabilize the SMO / PKA-C complex at the membrane inner leaflet.

Lastly, HDX-MS measurements revealed that PKA-C binding causes conformational changes in several regions of SMO that do not directly contact the kinase, including transmembrane helices 5 and 6 **(Extended Data Fig. 12a**), as well as its sterol-binding extracellular cysteine-rich domain (CRD)^6–11^ **(Extended Data Fig. 12b**). These conformational changes are reminiscent of the allosteric structural effects that occur when canonical GPCRs bind their cytoplasmic signaling partners^98–102^.

Overall, our findings illustrate how the SMO / PKA-C complex, despite an entirely different molecular composition, mimics the protein and lipid interfaces, as well as the allosteric conformational changes, seen in canonical GPCR assemblies with their effectors.

### A model for transmission of Hh signals by the SMO / PKA-C complex

Based on our findings, we propose the following model for how Hh signals are transduced across the membrane by the SMO / PKA-C complex (**Fig. 7**).

**Fig. 7:**
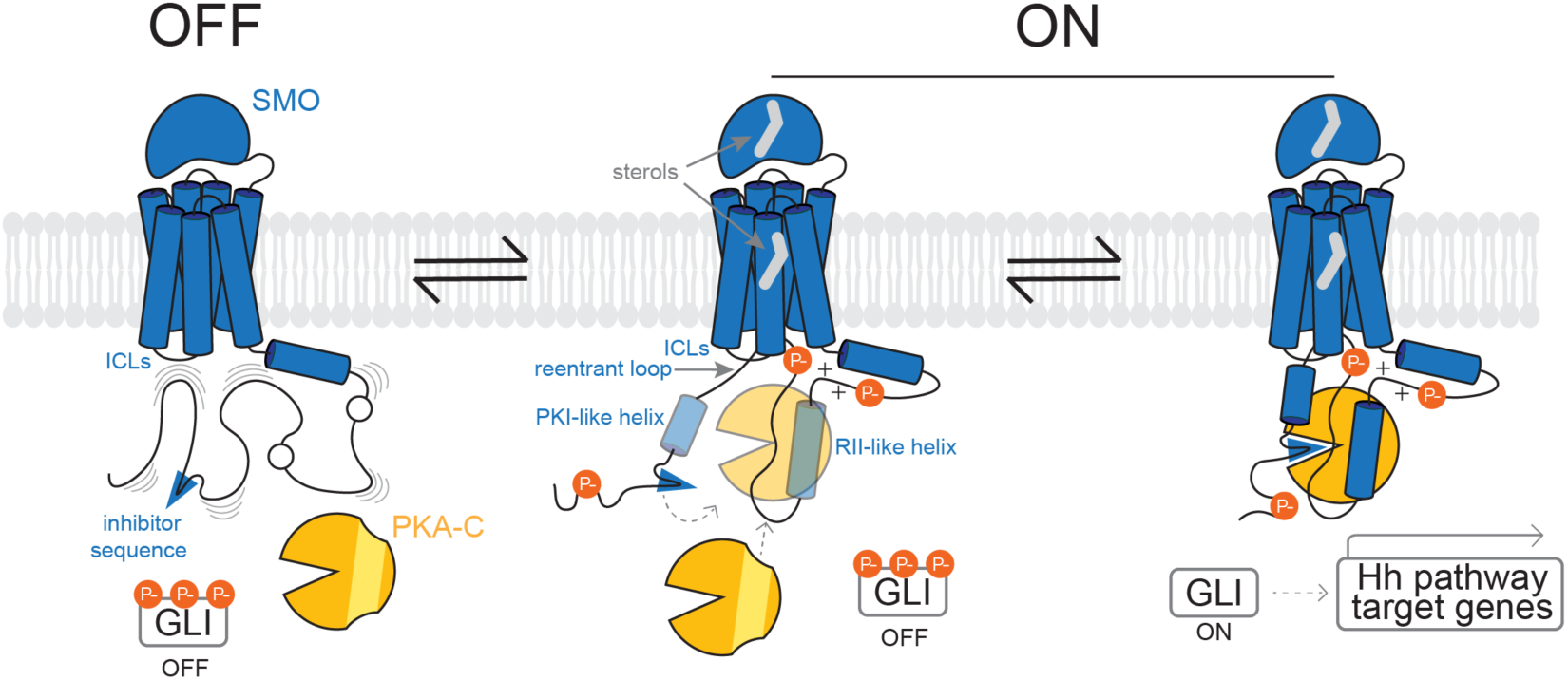
A structural mechanism for Hh signal transduction by the SMO / PKA-C complex. **(Left)** In the Hh pathway “off” state, SMO (blue) is inactive, with its pCT largely disordered (indicated by wavy lines) and unable to effectively engage PKA-C (orange) due to insufficient phosphorylation by GRK2/3. **(Middle)** In the Hh pathway “on” state, inhibition of PTCH1 (not shown) enables sterols (gray) to bind the SMO extracellular and 7TM domains, leading to SMO activation and phosphorylation by GRK2/3. This stabilizes secondary structures (PKI-like and RII-like helices) in the pCT and enables SMO to form a complex with PKA-C. PKA-C binding reinforces the SMO pCT secondary structures, further enhancing complex formation. **(Right)** The SMO inhibitor sequence (blue wedge) enters the PKA-C active site to interrupt PKA-C’s catalytic cycle. Consequently, GLI is released from phosphorylation-induced inhibition, leading to transcription of Hh pathway target genes. See main text for additional details.

In the pathway “off” state, SMO is inactive. As a result, the pCT is not efficiently phosphorylated by GRK2/3, remains largely disordered, and cannot effectively engage PKA-C (**Fig. 7, left panel)**.

In the pathway “on” state, Hh proteins bind to and inactivate PTCH1. Consequently, SMO binds membrane sterols^6–12^ and assumes an active conformation^11^, enabling recognition and phosphorylation by GRK2/3^13,15^. SMO phosphorylation stabilizes the pCT via intramolecular electrostatic contacts, enabling formation of secondary structures such as the RII-like helix and the PKI-like helix that mimic PKA-C-interacting regions in classical PKA inhibitors. Binding of PKA-C provides positive feedback to reinforce these structures, similar to the ‘folding upon binding” that occurs when other IDRs engage their specific targets^34,35^, such as when intrinsically disordered PKI proteins interact with PKA-C^29^. The SMO 7TM cavity stabilizes the SMO/PKA-C complex by bringing the PKI-like and RII-like helices closer together, while the SMO ICLs and membrane lipids provide additional PKA-C docking surfaces that enhance SMO / PKA-C complex formation via an avidity effect (**Fig. 7, middle panel)**. These interactions cooperate to facilitate insertion of the SMO inhibitor sequence into PKA-C’s active site which interrupts the enzyme’s catalytic cycle, inhibiting GLI phosphorylation to ultimately elicit transcription of Hh pathway target genes (**Fig. 7, right panel).**

Thus, SMO active state-dependent phosphorylation remodels SMO into a “parallel holoenzyme”, analogous to the PKA-R / PKA-C holoenzyme, that blocks PKA-C activity using principles borrowed from canonical GPCR and kinase signaling assemblies.

## DISCUSSION

By merging computational structural approaches with HDX-MS, biochemical studies, and functional assays, we have delineated a structural mechanism for a key step in Hh signal transduction, namely how the active state of SMO directly binds and inhibits PKA-C to transmit Hh signals intracellularly. Our study was enabled by AlphaFold^44,45,49^, which provide here the first structural snapshots of SMO’s intrinsically disordered C-terminus in its conditionally folded, PKA-C-bound conformation. The AlphaFold model aligns well with our computational and experimental findings, indicating that it depicts the SMO / PKA-C complex in a physiological state. Our work highlights the power of AlphaFold and similar algorithms to probe protein assemblies that have eluded empirical structural approaches. The strategy utilized here may be more broadly applicable to conditionally folding IDRs and has the potential to reveal the functions of this ubiquitous, essential, and poorly understood class of sequences^34–36,103^.

Our work reveals surprisingly extensive parallels between the SMO / PKA-C complex and canonical GPCR and PKA signaling assemblies. Like PKA-R subunits, SMO binds PKA-C by combining a central active site-binding inhibitor sequence with auxiliary elements contacting peripheral kinase surfaces^2,25,27^. This mimicry is exemplified by the SMO RII-like helix, which, in a remarkable example of evolutionary convergence, utilizes nearly the same strategy to engage PKA-C as does the PKA-RIIβ β4-β5 loop^32^, despite no discernible primary sequence or secondary structural homology. In the same vein, the SMO / PKA-C interaction echoes the tripartite binding mode of canonical GPCR / β-arrestin complexes, which utilize the receptor C-terminus, intracellular 7TM surface, and membrane lipids^16,17^.

However, a notable distinction between SMO and canonical GPCR signaling complexes lies in how phosphorylation triggers receptor / effector binding: while GPCR / β-arrestin complexes rely on insertion of a phosphorylated receptor peptide into a preformed, positively charged β-arrestin groove^70,76,104^, SMO phosphorylation induces a large-scale rearrangement of the SMO C-terminus into a partially ordered state defined by PKA-C-interacting structural elements. This mechanism is well-suited to regulate the SMO / PKA-C interaction, as it couples SMO activation to PKA-C binding while also taking advantage of well-established docking motifs on PKA-C, including the αF-αG region, ATP-binding pocket and αG-αH region^31,32,51^. The GRK2/3-phosphorylated SMO residues may directly stabilize the RII-like helix, as suggested by their proximity to conserved and essential K/R residues in or near this helix. Alternatively, or in addition, interactions between GRK2/3-phosphorylation sites and K/R residues may facilitate the folding of the SMO pCT by stabilizing key intermediates in the folding process, and thereby overcoming the energy barrier to achieving a folded state, similar to other proteins with phosphorylated IDRs^34,105,106^. In the Hh pathway “off” state, negatively charged lipids may anchor SMO pCT K/R residues at the membrane inner leaflet (as suggested by prior MD simulations^107,108^); this may inhibit SMO / PKA-C interactions by preventing spontaneous formation of SMO pCT secondary structures when SMO is not phosphorylated, ensuring tight control of Hh signal transduction when SMO is inactive.

Our AlphaFold prediction shows high pLDDT scores particularly in the RII-like helix and pseudosubstrate motif, indicating a high level of confidence in these regions of the model. The lower pLDDT scores in other SMO pCT regions may not arise from a deficiency in the AlphaFold prediction *per se,* but rather may reflect the lack of a single, stable conformation even in the presence of PKA-C. Such conformational heterogeneity is not unique to SMO, and is indeed characteristic of many GPCRs^16,17,82^, potentially explaining why large segments of GPCR intracellular domains remain disordered (and therefore unresolved) in empirical structures of GPCR-effector complexes. AlphaFold generated one major class of models for the SMO / PKA-C complex as well as a second, less prevalent class. These two classes may represent intermediates in the SMO-PKA signaling process, or unique signaling states that lead to distinct downstream outcomes. Future studies can address these possibilities and might reveal additional conformations of the SMO / PKA-C complex not captured by our models.

In conclusion, our work provides a structural mechanism for how SMO binds and inhibits PKA-C, illuminating a pivotal step in Hh signal transduction. Blocking the SMO / PKA-C interfaces characterized here may provide a new therapeutic strategy to thwart ectopic Hh signaling in cancers, especially when resistance to SMO orthosteric site inhibitors develops^109,110^. Beyond SMO, our study has broad implications for kinases and GPCRs. First, how PKA-C recognizes its physiological substrates remains structurally unresolved, as existing structures are mostly limited to PKA-C complexes with short peptide substrates^26,50,111,112^. Our study suggests that substrate IDRs, where most kinase phosphorylation motifs reside^113–115^, facilitate substrate recognition by forming transient PKA-C-interacting elements similar to the IDR in SMO, a hypothesis readily testable using the strategies described here. Second, numerous canonical GPCRs undergo dual phosphorylation by PKA and GRK kinases^116–118^, profoundly affecting receptor-mediated signaling, but how GRK phosphorylation influences PKA phosphorylation (and vice versa) remains poorly understood. The SMO / PKA-C complex may provide a valuable model to study the interplay between GRK and PKA binding to GPCRs at a structural level, shedding light on a widespread GPCR-regulatory mechanism. Finally, our work shows how GRK-mediated phosphorylation can induce large-scale remodeling and conditional folding of a GPCR’s IDR segments, enabling binding and regulation of an essential downstream effector. Phosphorylation-induced remodeling of IDRs may operate broadly throughout the GPCR superfamily to enable interactions with conventional^119^ or unconventional effectors, leading to an expanded array of regulatory complexes, downstream signaling outputs, and biological outcomes. Understanding and controlling these processes across GPCRs represents an exciting future challenge.

## Supporting information

Supplementary Data 1: AlphaFold 3 Models of SMO-PKA Complexes

Supplementary Data 2: AlphaFold 2.3 Models of SMO-PKA Complexes

## SUPPLEMENTARY INFORMATION

### Supplementary Data Files

Supplementary Data File 1: AlphaFold 3 models of SMO / PKA-C complex Supplementary Data File 2: AlphaFold 2.3.0 models of SMO / PKA-C complex.

### Supplementary Discussion 1 - 4

1. <u>AlphaFold predictions of IDRs in conditionally folded states:</u> AlphaFold tends to predict conditionally folded IDRs in their conditionally folded states, even when the post-translational modifications and/or interacting partners necessary for conditional folding are not included in the model. This is because the sequence-structure relationships captured by AlphaFold’s multiple sequence alignments reflect contributions from these factors, and therefore inform AlphaFold’s final outcome even when they are not explicitly modeled^46,47^. Thus, although AlphaFold 2.3 does not permit specification of phosphorylation sites, the SMO / PKA-C models appear to capture phosphorylated, conditionally folded forms of SMO. In support of this: (a) modeling SMO / PKA-C interactions with AlphaFold3, which enables phosphorylation, produced models that are very similar to those generated by AlphaFold 2.3.0 (**Fig. 1a, Extended Data Fig. 1c**); (b) the models agree closely with our HDX-MS, disulfide trapping, SPR, and functional studies of phosphorylated SMO / PKA-C complexes (**Fig. 3, 6**); (c) several PKA-C-binding secondary structural elements in the SMO pCT (e.g, pseudosubstrate motif, RII-like helix) are visible even in models of SMO alone. i.e., lacking PKA-C **(Extended Data Fig. 1b**). These observations underscore AlphaFold’s propensity to predict IDRs in their conditionally folded states.
2. <u>Two SMO / PKA-C models produced by AlphaFold 2.3.0:</u> AlphaFold 2.3.0 produces two classes of models for the SMO / PKA-C complex, which we term conformation 1 (conf1) and conformation 2 (conf2) **(Extended Data Fig. 1c**). Alignment of the two models on their PKA-C subunit reveals that the SMO pCT / PKA-C module is rotated with respect to the SMO 7TM domain. As a result, the SMO pCT, including the pseudosubstrate motif, RII-like helix, and reentrant loop, are in near-superimposable configurations in both models, while the SMO ICLs engage one of two distinct surfaces of PKA-C in each model. Both models have high ipTM scores (79-84 for conf1, 80-81 for conf2) and are consistent with our HDX-MS studies. While we primarily refer to the AlphaFold 3 model (similar to conf1) in our manuscript, conf2 may also be physiologically relevant. Intriguingly, conformational heterogeneity is also seen in GPCR-arrestin complexes, where the arrestin can rotate, presenting different surfaces to the GPCR 7TM domain ^17^.
3. <u>Disulfide trapping of SMO / PKA-C complexes for HDX-MS, and analysis of SMO</u> <u>pseudosubstrate motif and PKA-C active site cleft</u>: To enable our HDX-MS studies, we utilized disulfide trapping to stabilize the SMO / PKA-C complex. This strategy was advantageous because it (a) prevented untoward complex dissociation, and (b) allowed us to trap the nonphosphorylated state of the complex, permitting explicit comparison to the phosphorylated state, and thereby revealing how phosphorylation affects this complex. When interpreting our HDX-MS data, however, it is important to ensure that the results do not simply reflect artifactual, nonspecific interactions that may arise from covalently linking two proteins together into a complex. This scenario is unlikely because (a) SMO / PKA-C interaction results in protection of specific SMO and PKA-C regions known to be essential for each protein’s functionality **(Extended Data Fig. 5a, b**); (b) protection depends on agonist-induced GRK2 phosphorylation, as we observed substantially less protection of disulfide-trapped SMO / PKA-C complexes bound to the SMO inverse agonist KAAD-cyclopamine (KAADcyc) and lacking phosphorylation (**Fig. 5d, Extended Data Fig. 9a, b**). In contrast, if the HDX-MS results were nonspecific, protection would be independent of SMO phosphorylation and not involve specific, functionally relevant regions of SMO and PKA-C. The SMO inhibitor sequence is protected upon binding PKA-C **(Extended Data Fig. 5e, f**) regardless of SMO phosphorylation, due to the disulfide trap (SMO L637C to PKA-C C199). However, under physiological conditions (in which SMO and PKA-C are not covalently linked), this interaction likely occurs in a SMO activity-dependent manner, i.e. only when SMO is phosphorylated. Interestingly, the PKA-C active site cleft and substrate-binding region (C helix, activation loop, P+1 loop) become more dynamic upon SMO binding **(Extended Data Fig. 5e, f**). This may be due to: (a) the SMO L637C – PKA-C C199 covalent bond trapping the kinase in a partially open conformation, as observed in disulfide-trapped PKA holoenzymes^59,60^, and (b) the elevated pH (8.0) used, which favors disulfide trapping^60^ but likely hinders kinase domain closure due to the deprotonation of histidine H87 within the active-site cleft^120^. Thus, while SMO undergoes protection, parts of PKA-C become more dynamic, consistent with our model.
4. <u>SPR studies of PKA-C interactions with wild-type and mutant SMO:</u> The SMO IFR→AAA mutation significantly impairs GLI transcriptional activation in cultured cells (**Fig. 4a**) and reduces SMO/PKA-C binding in the in vitro SPR assay (**Fig. 4c, Extended Data Fig. 7**), supporting the idea that this mutation disrupts the SMO RII-like helix/PKA-C hinge interface. The impact of this mutation is less pronounced *in vitro* than *in vivo*, possibly due to: (1) a higher sensitivity of the *in vivo* GLI transcriptional assay to changes in amounts of the SMO / PKA-C complex over a relatively narrow range; (2) a greater extent of GRK2/3 phosphorylation of SMO in our *in vitro* SPR assay compared to our cell-based transcriptional activity assays^15^, which might stabilize the SMO / PKA-C complex and thereby blunt the deleterious effects of a mutation. Nevertheless, the weaker interaction of the SMO IFR→AAA mutant with PKA-C is consistent with our hypothesis that the RII-like helix participates in SMO / PKA-C interactions. We also note the higher affinity observed for the SMO / PKA-C complex in this study (K_D_ = 82 nM) compared to our previous measurements on PKA-C interactions with a soluble, nonphosphorylated SMO pCT fragment (K_D_ = 752 nM)^14^; we attribute this to the use of phosphorylated, near-full-length SMO and a membrane-like environment (nanodiscs) in the present study, reflecting a more physiological state of the complex.

## EXTENDED DATA FIGURES AND TABLES

**Extended Data Fig. 1:**
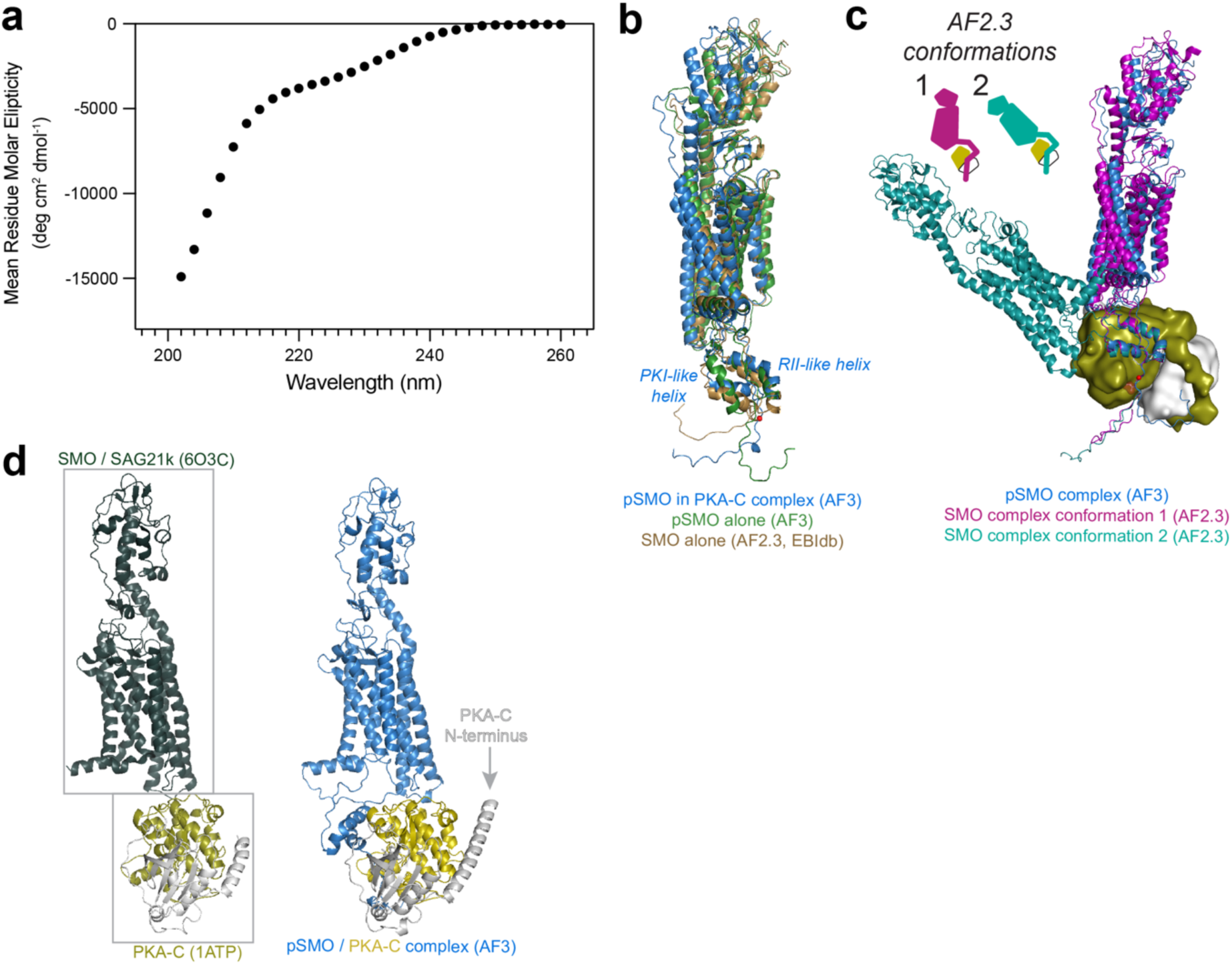
SMO binds PKA-C via its intrinsically disordered C-terminus. **a**, CD analysis of the purified SMO pCT, spanning residues 565-657 (40 μM). **b,** AlphaFold 3 models of SMO from the phosphorylated SMO / PKA-C complex (blue) aligned with the AlphaFold 3 model of SMO alone, with phosphorylation sites included (green), or the AlphaFold 2.3.0 model of SMO alone, downloaded from the EBI protein structure database^45^, in which phosphorylation sites were not explicitly specified (brown). **c,** Comparison of different SMO / PKA-C conformational states captured by AlphaFold 3 (blue) vs AlphaFold 2.3.0 (magenta or teal for conformations 1 or 2 (Conf1 or Conf2), respectively) (see **Supplementary Discussion 2**). Conf1 and Conf2 display near-identical positions for the PKI-like helix, RII-like helix, and reentrant loop, but with the PKA-C rotated such that it presents a new surface to the SMO 7TM domain. ipTM scores for Conf1 and Conf2 are 0.84 and 0.80, respectively. **d,** Left: empirical structures of agonist (SAG21k)-bound, active SMO (PDB: 6O3C) and PKA-C (PDB: 1ATP). Right: AlphaFold 3 model of the SMO / PKA-C complex. Note that in the AlphaFold model, the PKA-C N-terminus is pointing upwards the membrane.

**Extended Data Fig. 2:**
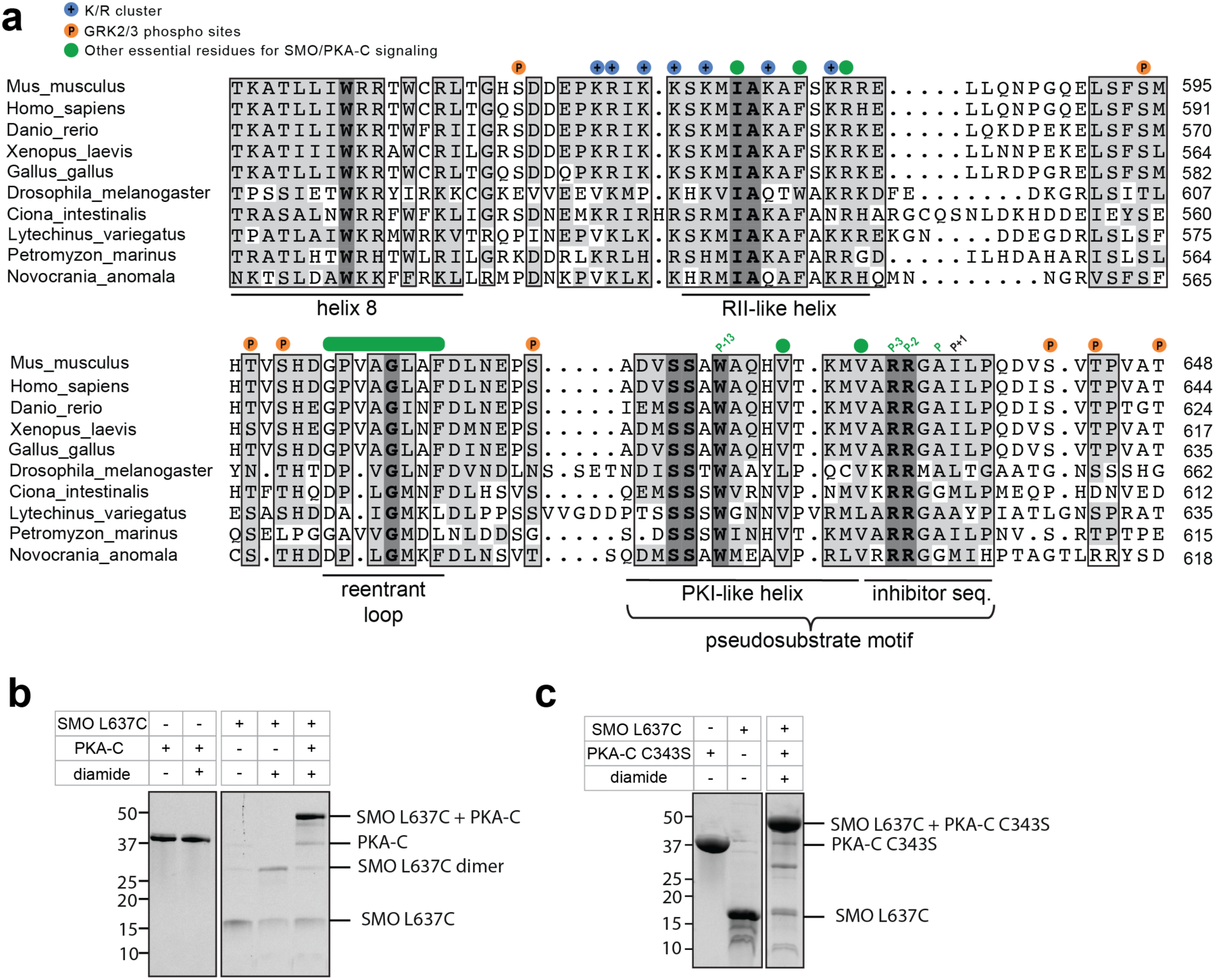
Evolutionary conservation of sequence and structural elements in SMO / PKA-C complexes. **a,** Multiple sequence alignment of helix 8 and pCT domains from SMO orthologs. Conserved K/R residues (blue circles), GRK2/3 phosphorylation sites (orange), and other essential residues for SMO / PKA-C signaling (green, see main text for detailed explanation) are indicated above the alignment, while secondary structural elements defined here or in prior studies are indicated below. **b,** Disulfide trapping of PKA-Cα with soluble SMO pCT L637C (i.e., lacking the CRD and 7TM domains), performed as in Fig. 2b. The SMO pCT has no endogenous cysteines, so trapping must be with L637C (the engineered Cys at P+2). **c,** Comparable disulfide trapping occurs with a PKA-C C343S mutant. Because PKA-Cα has only two endogenous cysteines (C199 and C343), this result indicates that disulfide trapping involves PKA-C C199.

**Extended Data Fig. 3:**
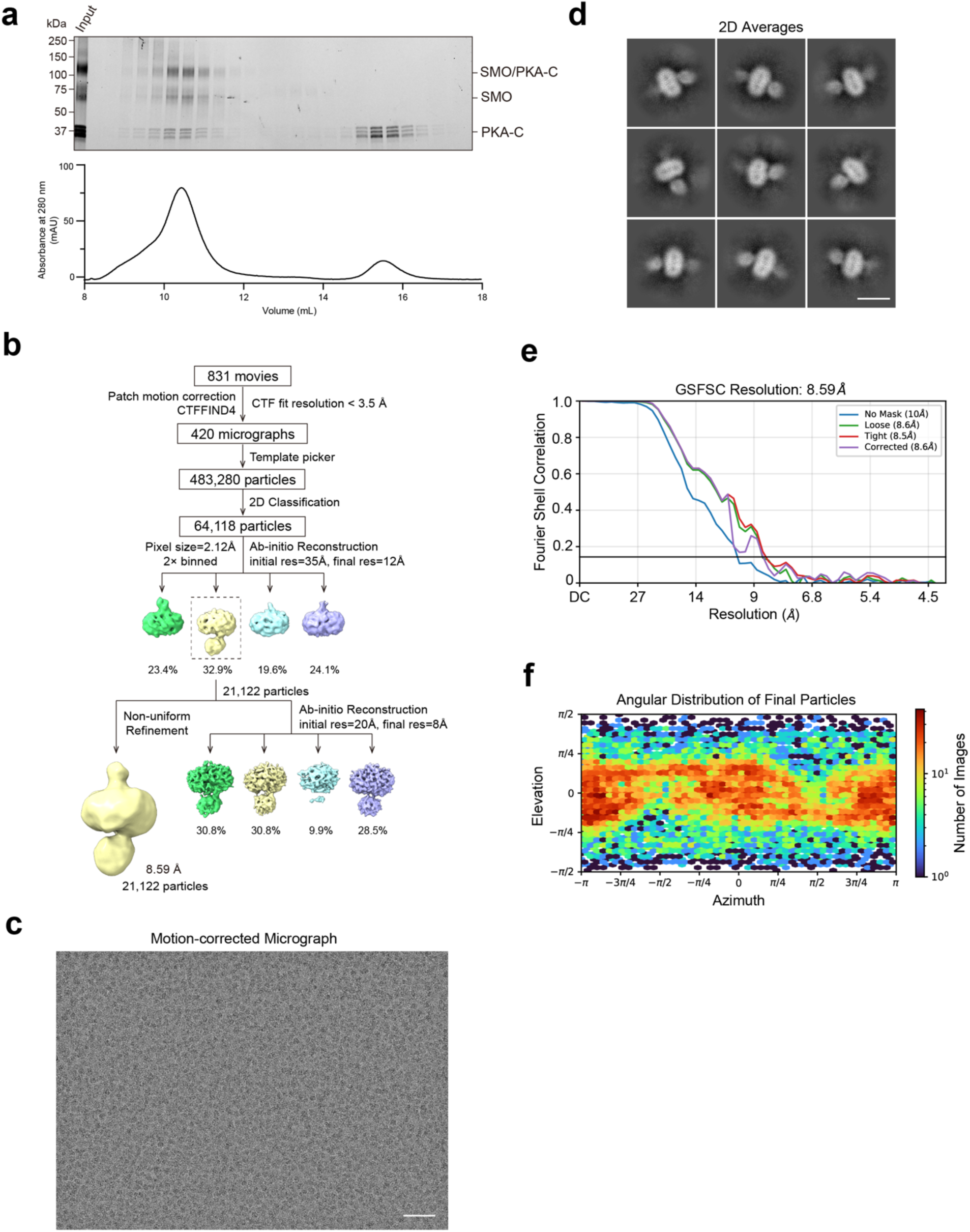
SMO/PKA-C complex preparation and cryoEM data processing. **a,** Size-exclusion chromatography and SDS-PAGE analysis of BS3-crosslinked SMO/PKA-C complex. **b,** Workflow of cryoEM data processing. **c,** Representative cryoEM micrograph (scale bar: 50 nm). **d,** Representative 2D class averages (scale bar: 10 nm). **e,** Gold-standard FSC curves of the EM map. **f,** Angular distribution plot of final particles.

**Extended Data Fig. 4:**
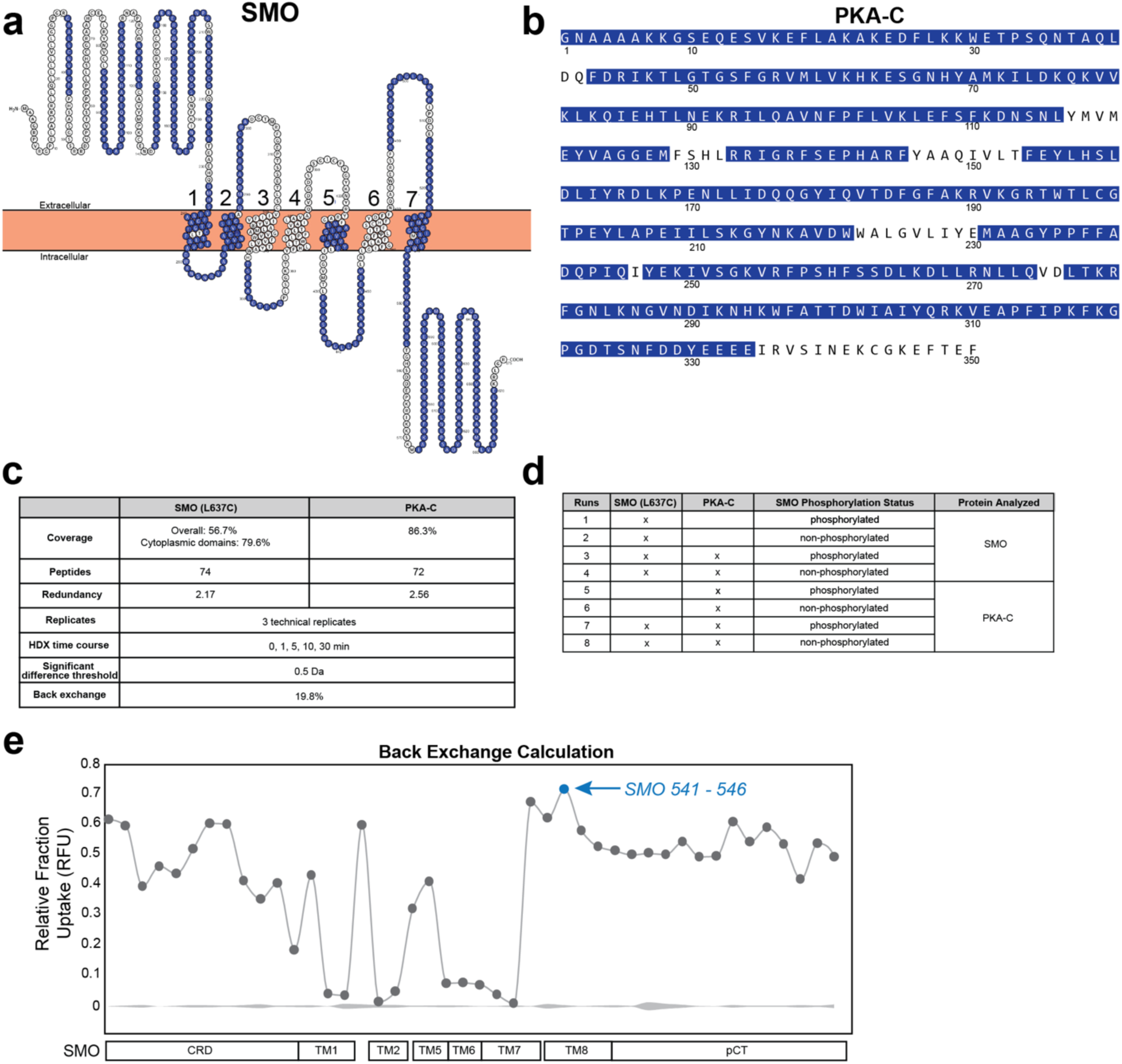
SMO and PKA-C sequence coverage in HDX-MS, and summary statistics of HDX-MS runs. **a,** Left: HDX-MS sequence coverage of the SMO L637C construct presented as a “snake plot”. Residues covered by the MS measurements are colored blue, while residues that were not detected by MS are colored white. TM helices are numbered in navy. **b,** HDX-MS sequence coverage of PKA-C. Colors are as in (**a**). **c,** Summary of protein coverage, # of peptides, and redundancy for MS runs. **d,** list of MS runs presented in this study (see Methods for details on sample preparation). **e,** Back-exchange for SMO 541-546, the most extensively deuterated peptide in our data set, was measured at 19.8% (see Methods).

**Extended Data Fig. 5:**
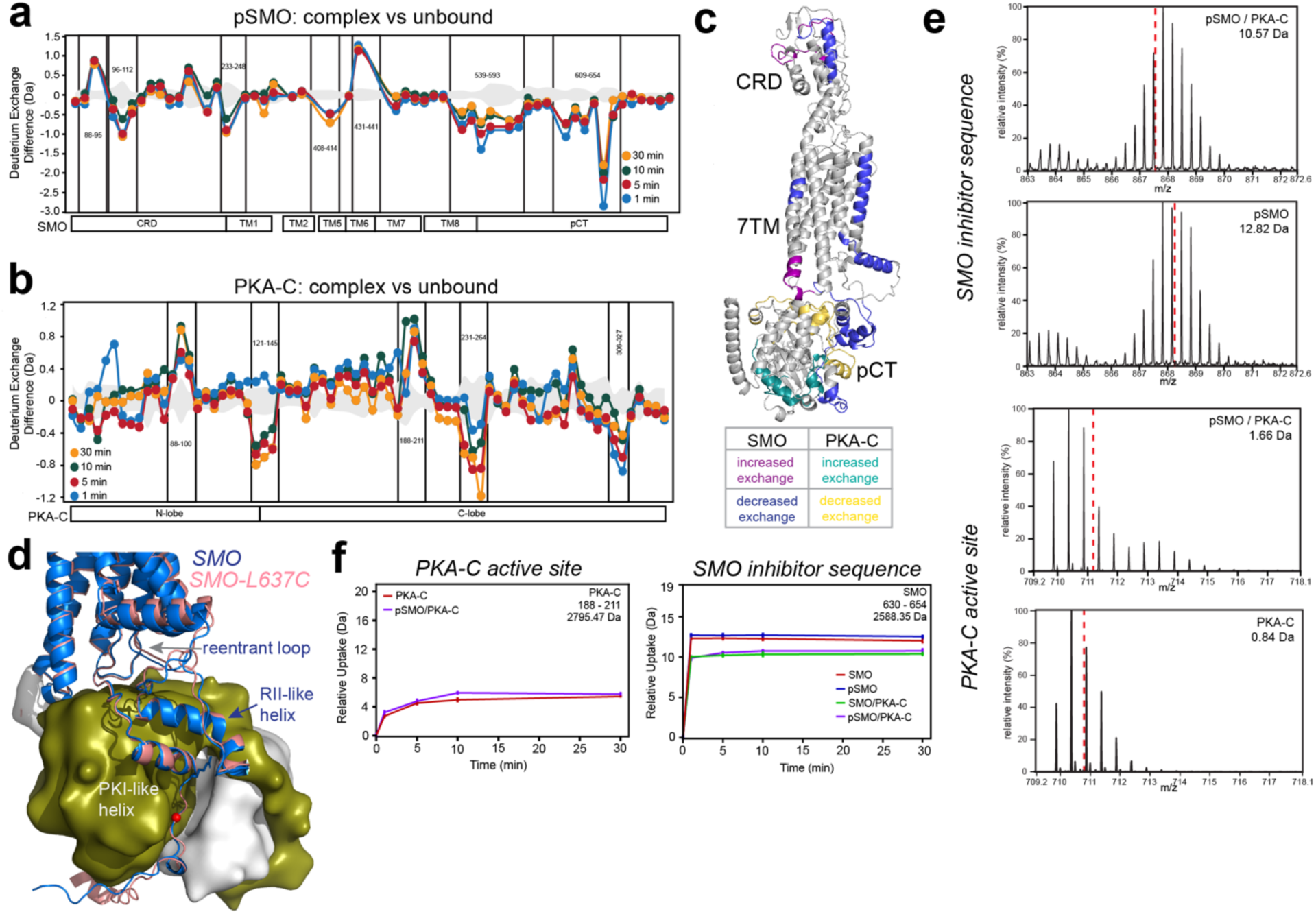
HDX-MS studies of phosphorylated SMO / PKA-C complexes vs. unbound proteins. Deuterium exchange difference (average number of deuterons) mapped for SMO (**a**) or PKA-C (**b**) in the phosphorylated SMO (pSMO) / PKA-C complex vs. the unbound protein, at the indicated timepoints following deuterium exchange. Key peptides are indicated on the plot, Negative differences denote decreased exchange (i.e., increased protection) in the complexed vs uncomplexed forms. Domain organization of SMO and PKA-C are indicated below each plot: CRD = cysteine rich domain, TM = transmembrane helix (numbered 1-7), pCT = membrane proximal C-tail. Standard deviations from replicate measurements are in gray. **c,** Mapping of HDX-MS results onto the AlphaFold3 model of the SMO / PKA-C complex. Peptides in SMO showing decreased or increased exchange (i.e, increased or decreased protection) are shown in blue and purple, respectively, while peptides in PKA-C showing decreased or increased exchange are shown in yellow and teal, respectively. **d,** Superposition of PKA-C complexes with wild-type SMO (blue) or SMO L637C (salmon), with key peripheral SMO structural elements indicated. P-site alanine is red. Although the PKI-like helix and RII-like helix are near-superimposable in both models, the reentrant loop is in slightly different positions within the SMO 7TM cavity; however, this is within the range of variability observed for this region even in the wild-type protein (see **Fig. 6d, Extended Data Fig. 10c**). Representative HDX-MS mass spectral envelope (t = 10 min) (**e**) or uptake vs. time plots (**f**) of the indicated peptides within the PKA-C active-site cleft or the SMO inhibitor sequence.

**Extended Data Fig. 6:**
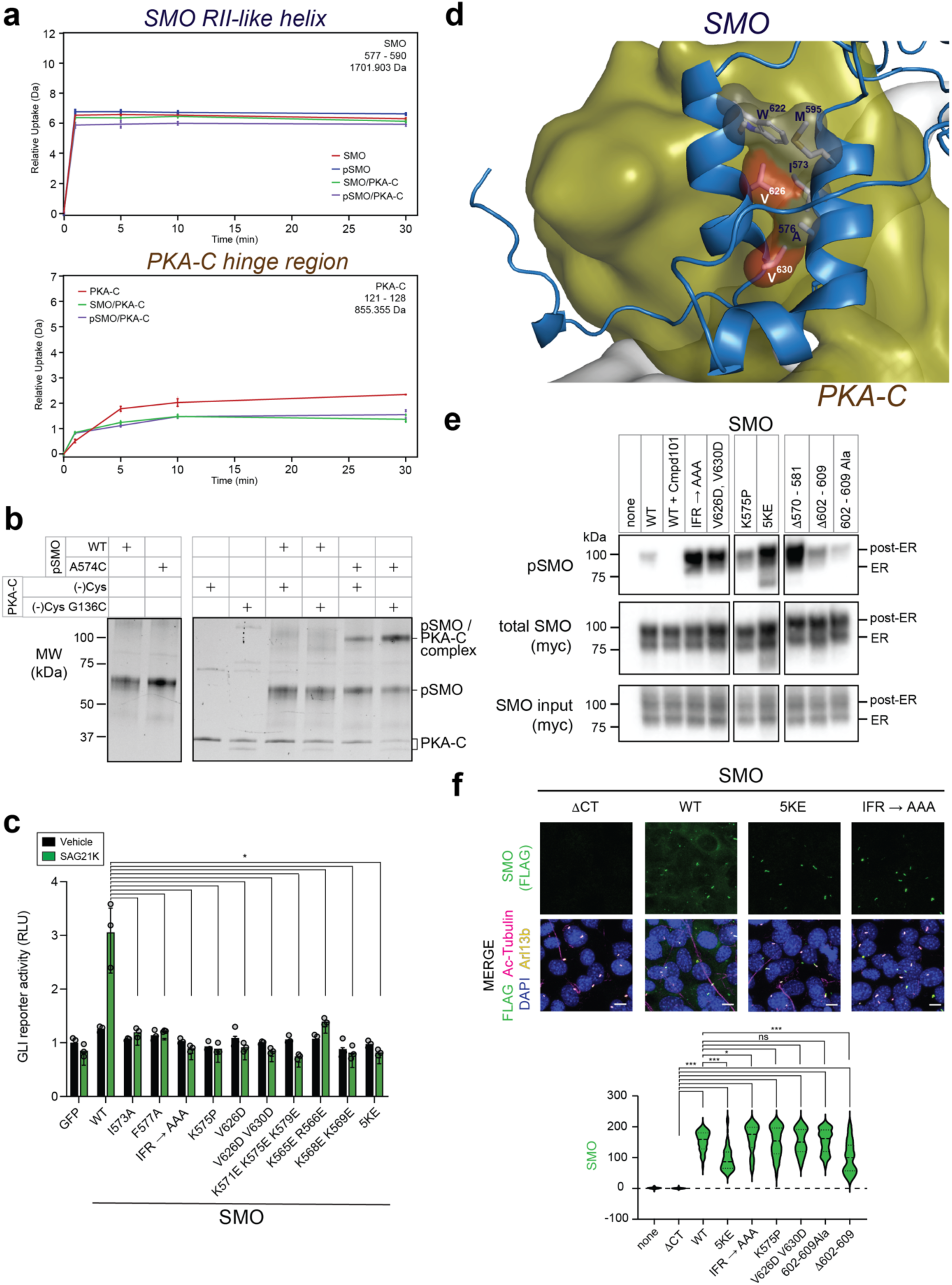
Additional biochemical, cell biological, and functional studies of the SMO RII-like helix. **a,** HDX-MS uptake vs. time plots for the indicated peptides in the SMO RII-like helix (top) and PKA-C hinge region (bottom). **b,** SDS-PAGE analysis of diamide-induced disulfide trapping between a minimal-cysteine PKA-C and SMO A574C, as in **Fig. 3e**. **c,** GLI transcriptional reporter assay was performed on the indicated SMO wild-type or mutant constructs following transfection into *Smo^-/-^* MEFs. The assay was conducted as in **Fig. 4a**, except using the direct SMO agonist SAG21k (green) vs a vehicle control (black). **d,** Locations of SMO V626 and V630 (red), and the residues with which they interact, at the interface between the PKI-like helix and the RII-like helix. **e,** SAG21k-induced phosphorylation of the indicated myc-tagged SMO wild-type and mutant constructs (or an empty vector control) was assessed following stable transfection into HEK293 cells, isolation of SMO-myc from cell lysates on anti-myc nanobody resin,and blotting with anti-pSMO antibody (top) and anti-myc antibody (bottom). Treatment of wild-type SMO with the GRK2/3 inhibitor Compound 101 (Cmpd101) serves as a negative control. Note that although several of these SMO mutants exhibit stronger phosphorylation than the wild-type protein, none of them exhibit less phosphorylation, making it unlikely that their deficits in GLI transcriptional reporter assays arise from an inability to undergo GRK2/3-mediated phosphorylation. **f,** Ciliary localization of the indicated FLAG-tagged SMO constructs was assessed following transfection into NIH3T3 cells, followed by SAG21k treatment (to induce SMO ciliary accumulation) and antibody staining. Left: representative images of selected SMO wild-type or mutant constructs, with FLAG-SMO (anti-FLAG, green), cilia (acetylated tubulin (AcTub), magenta; Arl13b, yellow) and nuclei (DAPI, blue). SMO lacking its entire C-terminus (SMOΔCT), which cannot localize to primary cilia^121^, serves as a negative control. Right: intensity-based quantification of FLAG-SMO localization to primary cilia. Note that although some of the mutants display a modest (<50%) decrease in ciliary localization compared to wild-type SMO, this is not sufficient to account for the near-complete loss of GLI activation in the transcriptional reporter assays. Scale bar = 10 microns.

**Extended Data Fig. 7:**
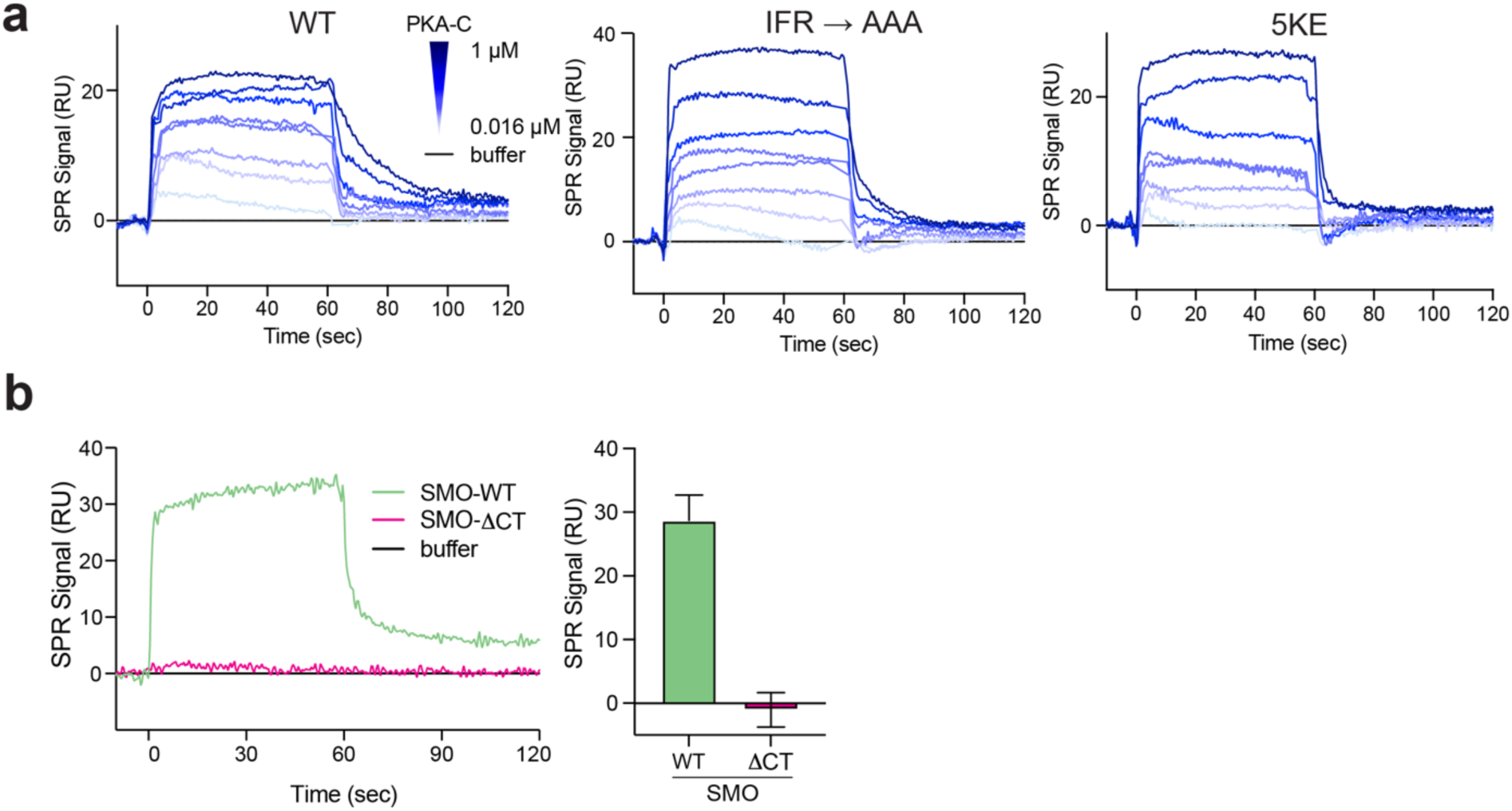
SPR studies of wild-type and mutant SMO / PKA-C complexes. **a,** SPR sensorgram for binding of myristylated PKA-Cɑ, in concentrations ranging from 0.016 μM to 1 μM, to phosphorylated wild-type (WT) SMO or the indicated SMO mutants reconstituted into nanodiscs. **b,** Left: SPR sensorgram for 250 nM PKA-Cɑ binding to wild-type SMO (green) but displaying no binding to a negative control version of SMO lacking the entire C-terminus (SMOΔCT, magenta), consistent with prior findings^13,15^. Right: Quantification of SMO WT or SMOΔCT SPR binding signal from n=3 replicates.

**Extended Data Fig. 8:**
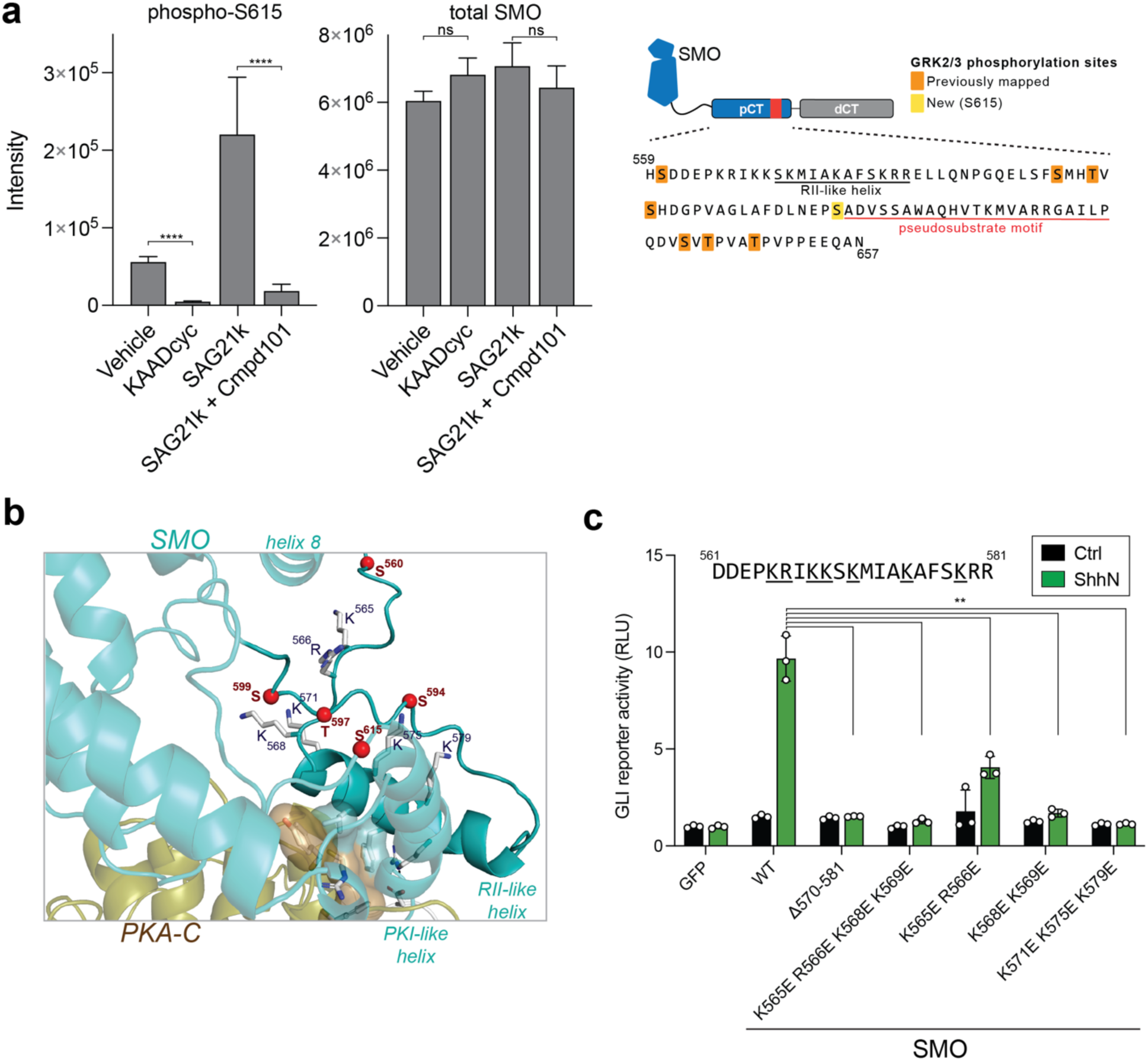
Additional biochemical and functional studies of SMO phosphorylation. **a,** Targeted MS-based quantification of phosphorylation of a phosphopeptide including pSMO 615 (left) and total SMO protein in each sample (middle), using FLAG-SMO protein isolated from HEK293 cells treated with vehicle, SMO agonist SAG21k, SMO inverse agonist KAADcyc, or GRK2/3 inhibitor Cmpd101. SMO sequence diagram indicating the location of S615 (yellow) along with the GRK2/3 phosphorylation sites previously mapped via MS^13^, is shown at right. “Intensity” is a measurement of the abundance of phosphorylation sites (left) or total protein (right), derived from model-based estimation in Msstats^122^ which combines individual peptide intensities. Inset depicts SMO pCT sequence with new (yellow) and previously mapped^13^ (orange) GRK2/3 phosphorylation sites. **b,** Location of GRK2/3-phosphorylated S/T residues and K/R residues in the alternative conformation of the SMO / PKA-C complex (AlphaFold 2.3.0 Conf2, see **Extended Data Fig. 1c**). **c,** GLI transcriptional reporter assay on *Smo^-/-^* MEFs transfected with the indicated SMO mutants and treated/analyzed as in **Fig. 5c**. SMO sequence from 561-581 is indicated, and residues mutated in this experiment are underlined.

**Extended Data Fig. 9:**
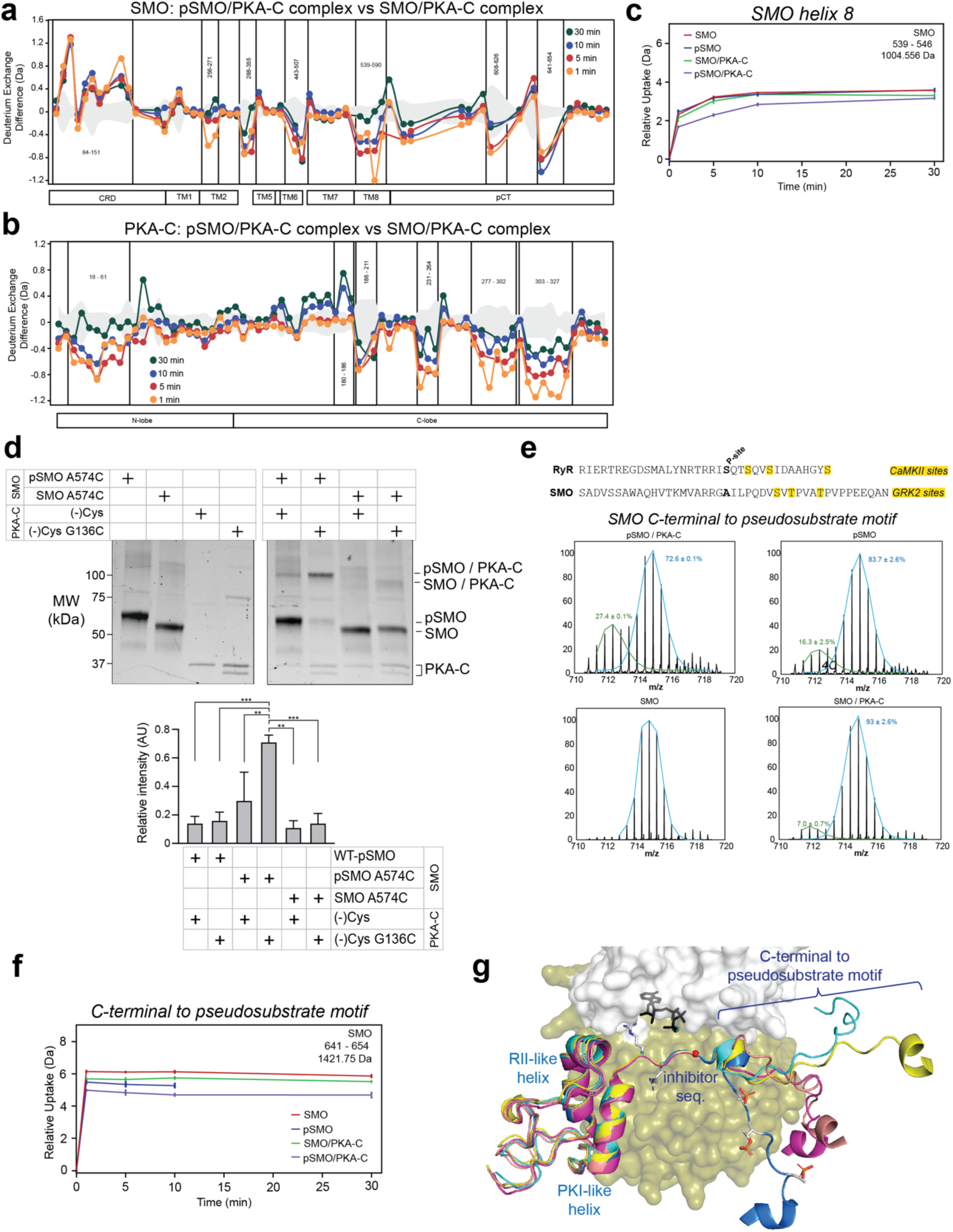
HDX-MS and disulfide trapping studies to monitor effects of phosphorylation on SMO / PKA-C complexes. Deuterium exchange difference (average number of deuterons) mapped for SMO (**a**) or PKA-C (**b**) in the phosphorylated vs nonphosphorylated SMO / PKA-C complex. Data are presented as in **Extended Data Fig. 5. c,** HDX-MS uptake vs. time plots for the indicated peptide in SMO helix 8. **d,** Disulfide trapping of SAG21k-bound, phosphorylated vs nonphosphorylated SMO A574C / PKA-C complex, performed as in **Fig. 5e**. Quantification is below the gel and represents the mean +/- s.d. from 3 replicates. **e,** HDX-MS uptake vs. time plots for the indicated peptide in the SMO region C-terminal to the pseudosubstrate motif. GRK2 phosphorylation sites in SMO, and CaMKII phosphorylation sites in RyR, are highlighted in yellow. **f,** HDX-MS uptake vs. time plots for the indicated peptide C-terminal to the pseudosubstrate motif (see **e**). Bimodal deconvolution of the peptide revealed high-exchanging (blue) and low-exchanging (green) populations, suggesting that this region may interact with PKA-C via a conformational selection mechanism. **g,** Overlay of top 5 AlphaFold 3 models indicating a propensity of the region C-terminal to the pseudosubstrate motif to wrap around PKA-C in some cases.

**Extended Data Fig. 10:**
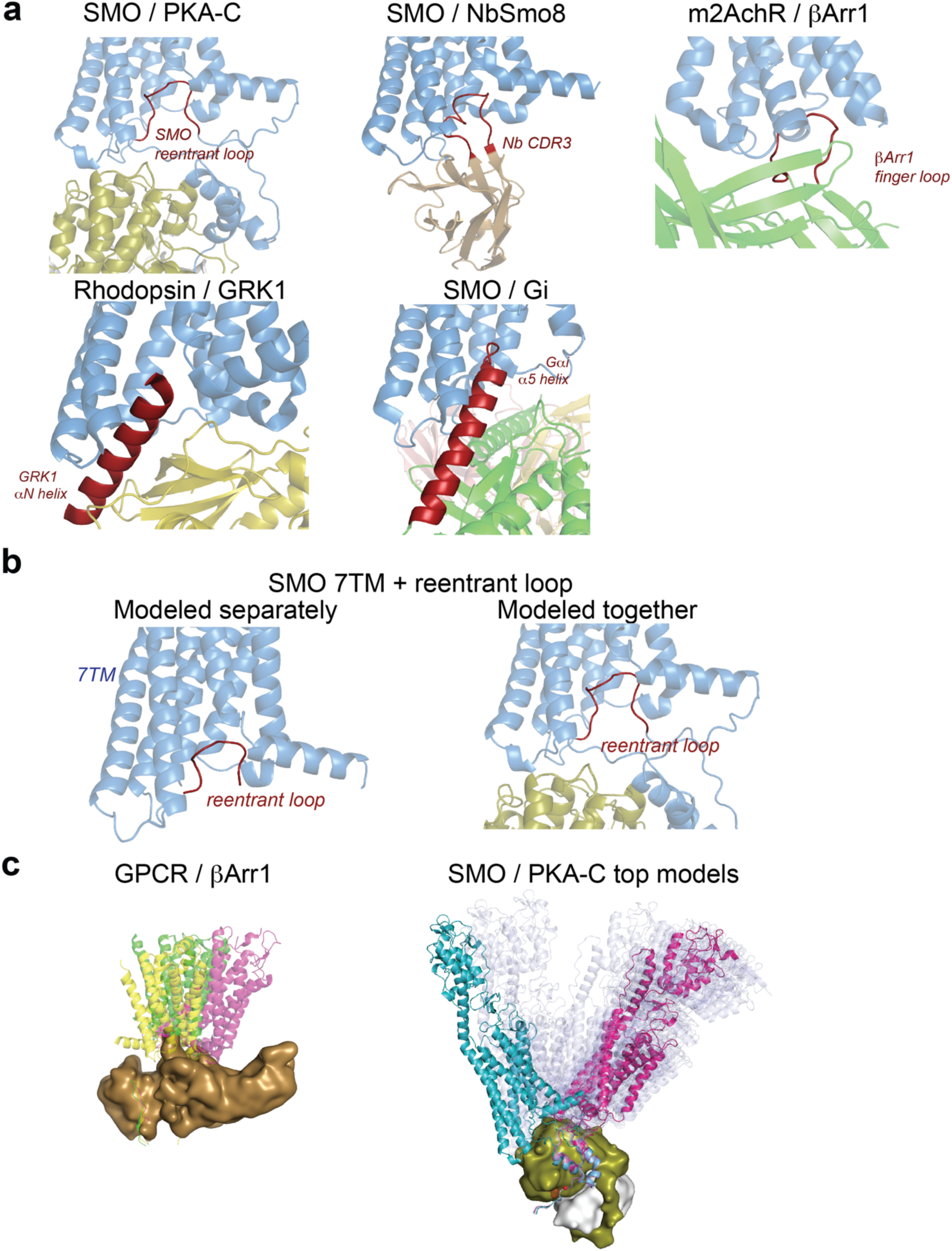
The SMO / PKA-C complex structurally mimics other GPCR-effector complexes. **a,** Comparison of the SMO / PKA-C complex to SMO or other GPCRs bound to the indicated proteins. In each structure, the GPCR portion is colored cyan, and the structural elements that engage the intracellular 7TM cavity are colored maroon. PDB numbers: SMO/NbSmo8 (6O3C), m2AchR/Barr1 (6U1N), Rhodopsin/GRK1 (7MT9), SMO/Gi (6XBL). **b,** Left: AlphaFold model of complex between a C-terminally truncated SMO construct (“7TM”) and the reentrant loop in the pCT, modeled as separate sequences. Right: AlphaFold model of near-full-length SMO (including the entire pCT) from the SMO / PKA-C complex, for comparison. The reentrant loop sequence is colored maroon in both models. **c,** Left: overlay of the following GPCR / β-arrestin structures, aligned on the β-arrestin: β1-adrenergic receptor /β-arrestin1 (6TKO), M2AchR / β-arrestin1 (6U1N), neurotensin receptor / β-arrestin1 (6UP7). Right: overlay of top 24 AlphaFold models of SMO / PKA-C complex (light blue), along with conf1 (pink) and conf2 (cyan) as shown in **Extended Data Fig. 1c**.

**Extended Data Fig. 11:**
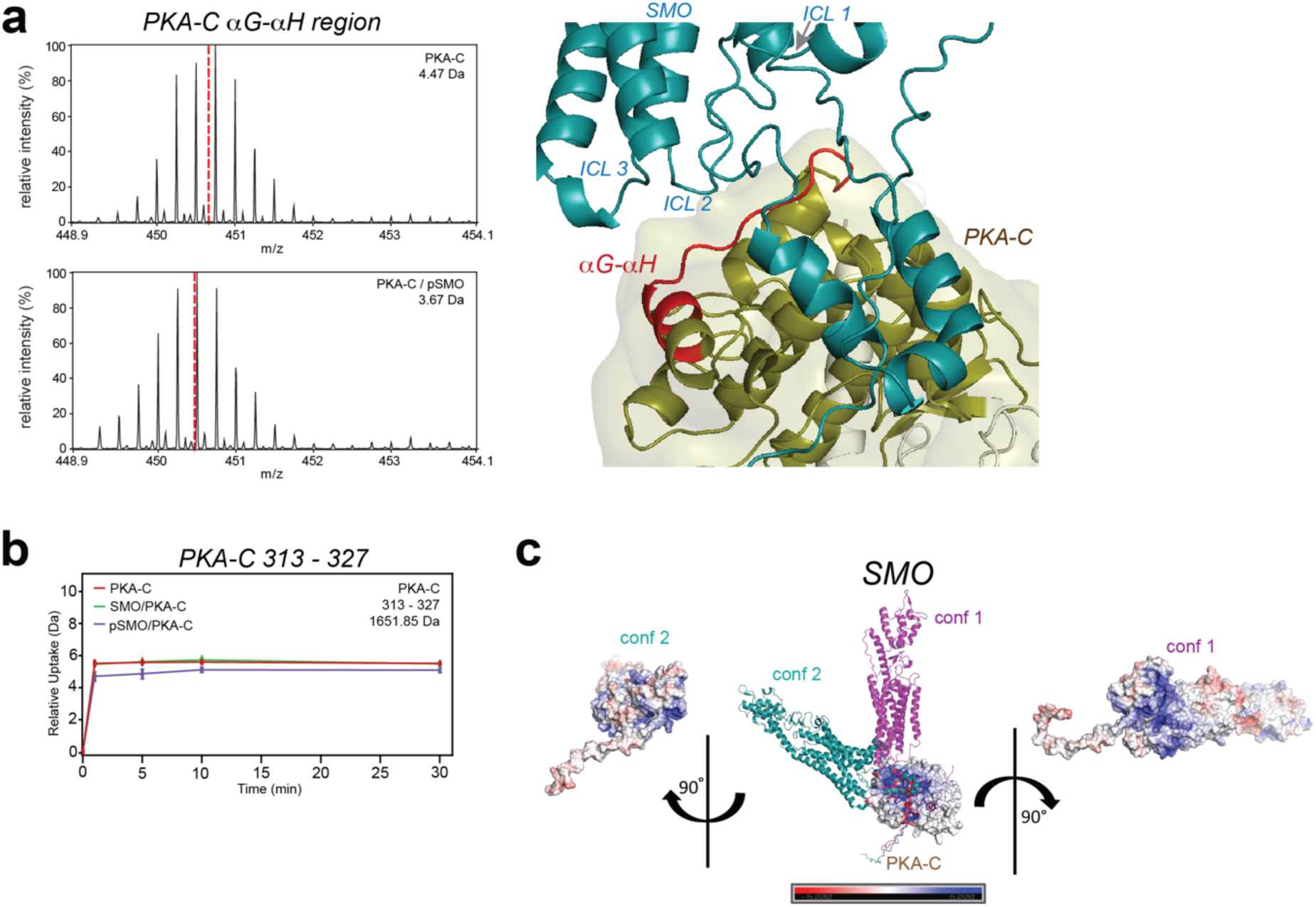
Influence of SMO ICLs and membrane lipids on SMO / PKA-C interactions. **a,** Left: representative HDX-MS mass spectral envelope (at t_ex_ = 5 min) for PKA-C 247-261 in the αG-αH loop. Right: αG-αH sequence protected in the SMO / PKA-C complex mapped onto conf2 of the SMO / PKA-C complex**. b,** Uptake vs. time plots of PKA-C 313-327. **c,** Electrostatic surface potential for each conformation (conf1, conf2) of the SMO / PKA-C complex. Scale is shown at bottom.

**Extended Data Fig. 12:**
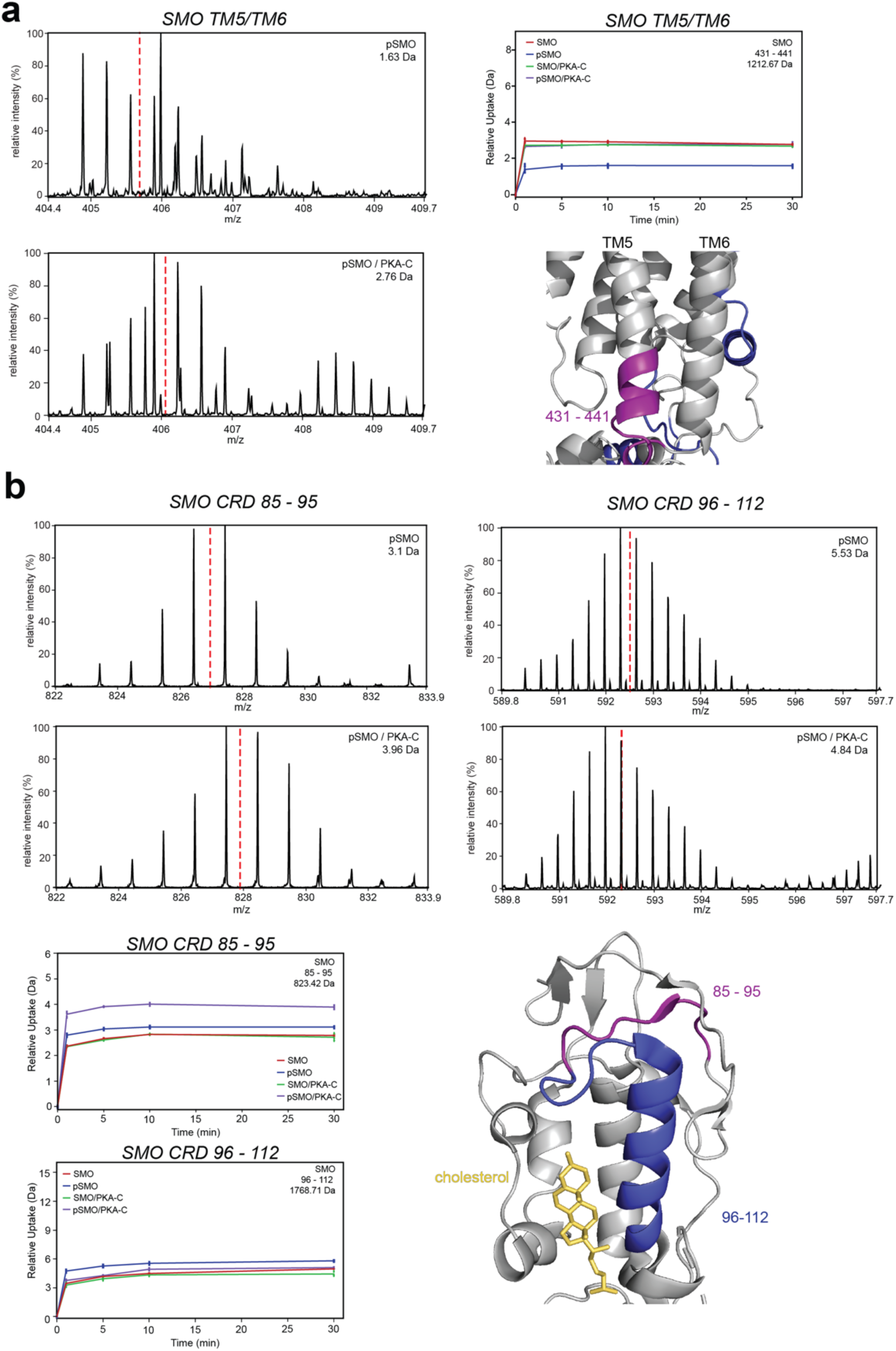
Allosteric changes in SMO induced by PKA-C binding. HDX-MS analysis of the indicated peptide in the TM5-TM6 region (**a**) or CRD (**b**) in phosphorylated SMO (pSMO) alone (top spectrum in each pair), or in complex with PKA-C (bottom spectrum in each pair). In each panel, representative HDX-MS mass spectral envelope and uptake vs. time plots are indicated at left, and regions of SMO that have undergone increased (purple) or decreased (dark blue) deuterium exchange upon SMO / PKA-C interaction are mapped onto the SMO-SAG21k-NbSmo8 structure (PDB: 6O3C)^11^.

## DATA AVAILABILITY

AlphaFold models of the SMO / PKA-C complexes have been deposited in the Zenodo database (see “Methods”). The molecular dynamics simulation trajectories have been deposited in GPCRmd (see “Methods”). Mass spectrometry data have been deposited at the Panorama server (see “Methods”). All unique biological materials are available upon request from the authors.

## METHODS

### AlphaFold modeling

AlphaFold 2.3.0 modeling was performed with localcolabfold^123^ downloaded from https://github.com/YoshitakaMo/localcolabfold on a Dell Alienware desktop machine with an RTX 4090 GPU of 24 Gbytes. We produced 50 models with 10 seeds and all 5 AlphaFold v2.3 weight sets without the use of templates from the Protein Data Bank (PDB). This was performed for various constructs (e.g. full-length mouse SMO and various truncations, and sequences from different species). A sample command is provided here: colabfold_batch --model-type alphafold2_multimer_v3 --zip --sort-queries-by none --amber --use-gpu-relax --num-seeds 10 --num-recycle 10 --recycle-early-stop-tolerance 1.0 smo_pka.fasta SMO/ > SMO.out Structures were ranked with the default function in Colabfold: 0.8*ipTM + 0.2*pTM. Structures were visualized in PyMOL (v.2.5, Schrödinger, Inc.).

AlphaFold 3 models were obtained from the AlphaFold3 server (https://golgi.sandbox.google.com/), which is the only form of AlphaFold3 available at this time. For models with PKA-C, two Mg^2+^ ions and ATP were included as ligands. Phosphorylation sites were manually entered via the PTM option on the server. Models were produced of complexes of phosphorylated mouse PKA-C (pT197 and pS338) and phosphorylated mouse SMO (GRK2/3 sites: pS560, pS594, pT597, pS599, pS642, pT644, pT648), along with non-GRK2/3 sites S578 and S666, all described previously^13^, as well as pS615, identified and described here (Extended Data Fig. 7a). Five random seeds were used (by submitting identical jobs 5 times) for a total of 25 models for each construct. The models were ranked by pairwise ipTM scores of PKA-C and SMO, with the top scoring model analyzed and discussed in Results.

As discussed above, AlphaFold 2.3.0 produced two distinct orientations of the GPCR domain of SMO with respect to the kinase domain of PKA-C, which we designated conf1 and conf2. The top ipTM scores of conf1 and conf2 were 0.84 and 0.80 respectively. AlphaFold3 models most closely resembled conf1 AlphaFold2.3 models. The highest scoring AlphaFold3 model had a chain_pair_ipTM of 0.78 for the SMO/PKA-C interaction.

For clarity the sequences of the highly disordered mouse SMO N-terminus (residues 1-64) and distal C-terminus (dCT) (residues 675-793) were removed from the structural models in the figure panels.

AlphaFold models discussed in this manuscript are available for download from Zenodo at the following URL: https://zenodo.org/records/13826713?token=eyJhbGciOiJIUzUxMiJ9.eyJpZCI6ImI0OWM4OGNmLWEwZWUtNDE5Mi1hYjUxLTExZTA3ZGM4ZDI0NSIsImRhdGEiOnt9LCJyYW5kb20iOiJmMmVlMmE4NjIwMzM4NzVjODYzMGE5Mjg2OWVmZTA2YSJ9.j6i3NOI5kjscJuGNtQ26KpWItN7CM-dAOoorGXMyWIxVEFAIxw2qNf-JdajyKPQkJ-iZfniZtvqvYmkEtja1mg

### Molecular biology

For SMO protein expression and purification, FLAG-tagged mouse SMO (residues 64-674, with N-terminal HA signal sequence, FLAG tags, and TEV protease site) or FLAG-SMOΔCT (as above, but with SMO residues 64-566) in pVLAD6 were described previously^13,15^. For expression and purification of the soluble SMO pCT, we used the pHTSHP vector as previously described^14^. For GLI reporter assays, full-length mouse SMO in the pGEN vector with C-terminal his and myc tags was described previously^6,13,14^. For ciliary localization studies, FLAG-SMO-nanoluc-IRES-mNG3k/pEF5-FRT-hygro was previously described^15^. N-terminally His-tagged GRK2 / pFastBac^124^ and mouse PKA-C in pRSET-b^14^ were previously described. A minimal-cysteine (Cys^-^) PKA-C construct (PKA-C C199A) was used as the background to introduce the PKA-C G137C mutation. Mutant DNA constructs in the above vectors were prepared in-house via Gibson assembly, or commercially (Epoch Life Sciences; Missouri City, TX), and verified by Sanger and/or next-generation sequencing before use.

### Cell culture and transfections

HEK293 suspension cells were cultured and transfected or BacMam-infected as previously described^13–15^. *Smo^-/-^* MEFs were cultured and transiently transfected as previously described^12–14^. NIH3T3 Flp-in cells were cultured and stably transfected via Flp-in integration, as previously described^15^.

### Small molecules, antibodies, and other reagents

SAG21k was obtained from BioTechne. Vismodegib was obtained from LC Laboratories (V-4050). KAADcyc was obtained from Toronto Research Chemicals (K171000). Cmpd101 was obtained from Hello Bio (HB2840). Control or ShhN conditioned medium was prepared as previously described^125^. The following antibodies were used in this study: mouse anti-FLAG M2 (Sigma, F3165), rat anti-Arl13b (BiCell Scientific, 90413), rabbit anti-AcTubulin (Enzo Life Sciences, BML-SA452-0100), rabbit anti-pSMO^15^ (7TM Antibodies, 7TM0239A), mouse anti-myc (clone 4A6, Millipore 05-724). Alexa-conjugated (Thermo Fisher) or HRP-conjugated (Promega) secondary antibodies were used for immunofluorescence or Western blotting detection, respectively.

### Protein purification and *in vitro* phosphorylation

To obtained phosphorylated SMO, FLAG-SMO was expressed in HEK293 Freestyle or HEK293 GnTI-cells, along with GRK2-eGFP, using the BacMam approach in the presence of 10mM sodium butyrate and 1 µM SAG21k and purified via FLAG affinity chromatography and gel filtration chromatography, then followed by in vitro GRK2 phosphorylation as previously described^15^. To obtain nonphosphorylated SMO, we followed the same procedure except that 1) SMO inverse agonist vismodegib was used in place of SMO agonist SAG21k during expression and purification; 2) the GRK2 coexpression and GRK2 *in vitro* phosphorylation steps were omitted, as previously described^15^. The soluble SMO pCT was expressed and purified from E. coli via sequential NiNTA affinity, SUMO tag cleavage / reverse NiNTA, cation exchange, and gel filtration as previously described^14^, with the following modifications: (1) *E. coli* were grown in LB rather than TB; (2) expression was induced by adding IPTG and incubating the cells at 30°C for 5 hrs; (3) the SUMO tag cleavage and overnight 4°C dialysis steps were combined. MSP1E3D1 scaffold protein was expressed and purified as previously described^126^.

His-tagged GRK2 was expressed in High Five cells via baculovirus and purified via NiNTA affinity chromatography and gel filtration chromatography as previously described^15,124^.

Wild-type, nonmyristylated mouse PKA-C was used for HDX-MS and disulfide trapping studies, and was expressed in *E. coli* and purified via IP20 chromatography as previously described^14,127^. SPR studies utilized myristylated PKA-C, which was prepared by coexpression with yeast N-myristyltransferase in *E.coli* BL21(DE3) cells as described previously^128^. A K7C mutation was introduced into PKA-C to increase myristylation efficiency in *E. coli*, as described previously^129,130^, and myristylated PKA-C was purified via IP20 chromatography. To confirm myristylation, IP20 purified PKA-C was run over a MonoS column and eluted protein samples were dialyzed into buffer containing 20mM KH2PO4 pH 6.5, 50mM KCl and 1mM TCEP and submitted to the Molecular Mass Spectrometry Facility at UC San Diego. Intact protein analysis was performed by using an Agilent 6230 time-of-flight mass spectrometer (TOFMS) coupled with an Agilent 1260 liquid chromatography (LC) system. The Jet Stream ESI source was operated under positive ion mode with the following parameters: VCap = 3500 V, fragmentor voltage = 175 V, drying gas temperature = 325 °C, sheath gas temperature = 325 °C, drying gas flow rate = 10 L / min, sheath gas flow rate = 10 L / min, and nebulizer pressure = 40 psi. The chromatographic separation was performed at room temperature on a Phenomenex Aeris Wide-pore C-4 column (2.1 mm ID x 50mm length, 3.6 µm particle size). Mobile phase A was HPLC-grade water with 0.1% TFA, and HPLC grade Acetonitrile with 0.1% TFA was used as mobile phase B. The mobile phase was delivered at a rate of 0.3 ml/min under gradient conditions as followed: Increased from 5% mobile phase B to 90% mobile phase B in 12 minutes, held at 90% mobile phase B for 2 minutes, returned to 5% mobile phase B in 1 minute, and equilibrated with 5% mobile phase B for 7 minutes. Agilent MassHunter software was used for data acquisition and analysis.

### SPR studies

For SPR studies, SMO was reconstituted into biotinylated MSP1E3D1 nanodiscs via a minor modification of previously published procedures^126^, summarized below. His-tagged MSP1E3D1 scaffold protein biotinylated as previously described^126^ except that a 1:10 molar ratio of MSPE3D1 to NHS-biotin was used, and the extent of biotinylation was assessed by monitoring capture of biotinylated MSPE3D1 on streptavidin magnetic beads (Thermo Fisher, 88816). SMO proteins were subject *in vitro* phosphorylation and purification as previously described^15^, except that we eliminated the gel filtration step following FLAG purification of the GRK2 phosphorylation reaction, and proceeded directly to nanodisc reconstitution. Nanodiscs were assembled using a 1:10:857:2571 molar ratio of SMO:MSP1E3D1:lipid:cholate in 1x HEPES-NaCl-EDTA (HNE) buffer. A lipid mixture consisting of 92 mol% 3:2 POPC:POPG and 8 mol% cholesterol (dissolved in chloroform) was placed in a borosilicate tube and dried under a stream of N2 gas to eliminate chloroform, followed by a 1-hour vacuum desiccation at room temperature. Subsequently, sodium cholate was introduced to the lipid mixture, followed by bath sonication for 10 minutes. Upon addition of water, an additional 5 minutes of sonication was performed to ensure complete lipid dissolution. Then, 20X HNE buffer (400 mM HEPES, 2000 mM NaCl, 20 mM EDTA), MSP1E3D1 protein, and SMO was added to the lipid mixture, and incubated for 3 hours at 4°C. Detergent removal was achieved by adding Bio-Beads (Bio-Rad) at a concentration of 14.5 mg per 100 μl reaction volume, followed by overnight rotation at 4°C. The reconstituted SMO was supplemented with 5 mM CaCl_2_ and applied to an M1 FLAG affinity column (to remove excess MSP1E3D1 and GRK2 leftover from the *in vitro* phosphorylation step). The sample was washed with 20 mM HEPES pH 7.5, 100 mM NaCl, and 5 mM CaCl_2_, then eluted using a buffer containing 20 mM HEPES pH 7.5, 100 mM NaCl, 1 mM EDTA, and 0.2 mg/mL FLAG peptide.

SPR interaction studies were performed in running buffer (20 mM MOPS, pH 7.0, 150 mM NaCl, 100 μM EDTA, 1 mM ATP, 10 mM MgCl_2_, 0.001% P20 surfactant) at 25 °C using Biacore 3000 instrument (GE Healthcare). Measurements were performed using a Biotin CAPture Kit (Cytiva) to capture biotinylated SMO Nanodiscs according to the manufacturer’s instructions. Briefly, Biotin CAPture reagent was immobilized to a level of 3,000 response units (RU) on the Sensor Chip CAP at the beginning of each cycle, followed by the sequential capture of respective SMO Nanodiscs (WT, DCT, 5KE, IFR→AAA) with a capture level of 250-350 RU on separate flow cells (flow rate 10 μl min^−1^). Serial dilutions of myr PKA-Cα K7C (16 nM – 1 μM) were diluted in running buffer and injected with increasing concentrations at a flow rate of 10 μl min^−1^ for 60 s (association) followed by 60 s dissociation in running buffer without PKA-Cα. Data were corrected (double referencing) for nonspecific binding and buffer effects by subtracting SPR signals from a flow cell with only CAPture reagent as well as blank runs by injecting buffer without the analyte using Biacore 3000 Evaluation Software 4.1.1 (Cytiva). The sensor chip was regenerated by three sequential injections of 6 M GuHCl/0.25 M NaOH, to remove the CAPture reagent until the baseline level was reached. Steady-state analysis was performed with GraphPad Prism 8.0.1 (GraphPad Software, Inc).

### Analytical-scale SMO / PKA-C disulfide trapping studies

Phosphorylated or nonphosphorylated forms of wild-type or mutant FLAG-SMO were prepared as described above, and gel filtered into 20 mM HEPES pH 7.5, 150 mM NaCl, 0.025% GDN, 1 μM SAG21K. PKA-C proteins were buffer exchanged into this same buffer using Zeba 7k desalting columns. To accurately compare disulfide trapping efficiencies between wild-type and mutant proteins, we quantified each protein’s concentration by A_280_ (adjusted for the extinction coefficient), then subjected each sample to a 2-fold set of serial dilutions which were analyzed via Stain-Free imaging of SDS-PAGE gels; the concentrations of each protein were then manually adjusted as necessary to ensure uniform concentrations between each set of wild-type and mutant proteins. For the reaction setup, 4 μL of SMO (12.5 μM), 4 μL of PKA (6.25 μM), 1 μL of 10X reaction buffer (10 μM SAG21K, 10 mM ATP, 100 mM MgCl_2_), and 1 μL of 50 mM NaOH (to adjust reaction pH to 8.0). Samples were then incubated for 30 minutes at room temperature to promote SMO / PKA-C complex formation. Following the initial incubation, diamide (prepared fresh before use) was added to each reaction at a final concentration of 5 mM to initiate disulfide bond formation, and samples were incubated for 1 hour at room temperature. Subsequently, 10 μL of 2X Laemmli sample buffer (without reducing agent) was added to each sample, which were then loaded onto a 4-20% SDS-PAGE gel. Disulfide trapped quantification was completed by analyzing adjusted, background-subtracted band intensities using Fiji. For each disulfide trapping condition, the adjusted intensity of the disulfide-bonded product was normalized by dividing it by the sum of the adjusted intensities of both the disulfide-bonded product and free SMO.

A similar protocol was employed for the disulfide trapping of soluble SMO pCT L637C with human PKA-C, with the following modifications. Both SMO pCT L637C and human PKA-C (wild-type or C343S) were buffer-exchanged into 20 mM HEPES pH 8 and 150 mM NaCl following gel filtration. During complex formation, 10 mM ATP and 100 mM MgCl_2_ were added, and the final concentrations of SMO pCT and PKA-C were 40 μM and 20 μM, respectively. The mixture was incubated for 3 hours following addition of 5 mM diamide, and analyzed via SDS-PAGE as described above for the near-full-length FLAG-SMO experiments.

### CD studies

SMO pCT(565-657) was purified as described previously^14^. Peak fractions from gel filtration containing intact SMO pCT were pooled and subjected to dialysis. Dialysis was performed using 3.5 kDa molecular weight cutoff tubing in 1x PBS (pH 7.4) at 4°C for two overnight cycles. Post-dialysis, the protein concentration was adjusted to 40 μM. Circular Dichroism (CD) measurements were conducted using an AVIV model 410 circular dichroism spectrometer. Samples were analyzed in a 1 mm path-length, quartz cuvette at 25°C, recording data every 2 nm across a wavelength range of 200-260 nm with a 3 s averaging time. Each condition, including a blanking control, was measured in quintuplicate. The collected raw data were averaged and blank-subtracted before normalization to mean residue molar ellipticity ([θ] = 100 * θ/(C * l * n), where C is concentration of protein in mM, l is path length in centimeters, and n is the number of peptide bonds in the protein) using a custom Python script. Data points with dynode voltages exceeding 500 V were excluded (specifically at the 200 nm wavelength).

### Preparative scale SMO / PKA-C disulfide trapping for HDX-MS

Large-scale pSMO L637C / PKA-C disulfide trapping for HDX-MS followed a similar procedure as described in the preceding section, except that SMO and PKA-C were present at 12 μM and 24μM, respectively, and the reaction was scaled up to 0.5 ml, and the diamide-induced disulfide bond formation proceeded for 2 hours. The reaction was then subjected to gel filtration chromatography on a Superdex 200 Increase 10/300 GL column equilibrated in 20 mM HEPES pH 8, 50 mM NaCl, 0.01% GDN, 1 μM SAG21k, 10 mM MgCl2, and 1 mM ATP. Fractions were analyzed by nonreducing SDS-PAGE, and those containing the pSMO / PKA-C complex were pooled and concentrated. The same procedure was used for disulfide trapping of nonphosphorylated SMO L637C / PKA-C; note that to obtain nonphosphorylated SMO samples, we took the following steps during purification (see “protein purification and *in vitro* phosphorylation” section above): vismodegib was used in place of SAG21k, GRK2 was not coexpressed with SMO, and purified SMO was not subjected to *in vitro* GRK2 phosphorylation.

### GLI reporter assays

GLI reporter assays on *Smo*^-/-^ MEFs transiently transfected with wild-type or mutant contructs, along with 8xGli-Firefly and SV40-Renilla dual luciferase reporter plasmids, were performed as previously described^12,13^.

### Monitoring phosphorylation of wild-type or mutant SMO via immunoblotting

HEK293 Freestyle cells were cultured in Freestyle 293 Expression Medium supplemented with 1% Fetal Bovine Serum, as previously described^12–15^. For transfection, 3 mL of HEK293 Freestyle cell culture (density: 3.0–4.0 million cells/mL) was transferred to a 6-well plate and incubated with 5 mM sodium butyrate at 37°C in a 5% CO_2_ atmosphere for 15 minutes. Transfection complexes were prepared by combining 1.25 μg of SMO WT or mutant DNA with 1.25 μg GRK2-GFP DNA in 250 μL OPTI-MEM I Reduced-Serum Medium. The mixture was vortexed, followed by the addition of 7.5 μL of TransIT-293 Reagent. After vortexing for 15 s, the mixture was incubated at 25°C for 15 minutes. The transfection complex was then added dropwise to the cell culture and incubated in a shaking incubator at 37°C and 5% CO_2_ for 48 hours. Cells were treated with either SAG21k or SAG21k + Cmpd101 for 4 hours, harvested by centrifugation at 2000 x g for 10 minutes, flash-frozen in liquid nitrogen, and stored at −80°C.

Cell pellets were resuspended in 500 μL of SDS-free RIPA buffer (50 mM Tris, pH 7.5, 150 mM NaCl, 2% NP40, 0.25% sodium deoxycholate, 10% glycerol) supplemented with Pierce protease and phosphatase inhibitor tablet. The samples were homogenized by pipetting and incubated at 4°C for 1 hour with rotation. Lysates were clarified by centrifugation at 21,100 x g for 10 minutes. An input sample was separated, and the remaining supernatant was incubated with 10 uL of ChromoTek Myc-Trap Magnetic Agarose at 4°C for 1 hour with rotation. The resin was washed three times with 1 mL of SDS-free RIPA buffer. Proteins were eluted by adding 50 μL of 2x Laemmli sample buffer containing β-mercaptoethanol and incubating at 25°C for 5 minutes.

Input and eluted samples were separated by SDS-PAGE using Criterion Stain-Free gels, transferred to PVDF membranes, blocked with 5% milk, and probed overnight with either rabbit anti-pSMO (7TM Antibodies, 7TM0239A) or mouse anti-myc (clone 4A6, Millipore 05-724). The relevant secondary antibodies were applied for 2 hours, followed by chemiluminescent detection using the ChemiDoc system (Bio-Rad).

### SMO ciliary localization studies

Ciliary localization studies on FLAG-tagged SMO stably expressed in NIH3T3 Flp-in cells were formed as previously described^15^. Briefly, stably transfected cells were grown to confluency on glass coverslips, switched to low-serum medium (DMEM + 0.5% FBS + pen-strep-glutamine) overnight to induce ciliation, then treated for 4 hr to induce SMO ciliary accumulation. Cells were washed, fixed in PFA, permeabilized with 0.1% Triton X-100, blocked overnight in TBST + 2% BSA. Cells were stained with anti-FLAG to identify stably expressed SMO and anti Arl13b and anti acetylated tubulin to mark cilia, followed by appropriate secondary antibodies + DAPI counterstain. Coverslips were mounted onto a slide with SlowFade mounting medium. Images were acquired on a Leica SP8 laser scanning confocal using a 40x water immersion lens. Identical zoom factors, exposure times, and gain settings were used in all experiments. SMO signal in cilia was quantified, background-subtracted, and graphed. Data for each condition represent 100 cilia counted from two or more separate fields in two independent trials.

### MD simulations

Systems for simulations were generated using CHARMM-GUI^131–133^ with parameters for the protein component assigned from the CHARMM36m forcefield^134^ and for other components from the CHARMM36 force field^135^. The AlphaFold 2.3.0 models of the SMO / PKA-C complex (conformation 1 or 2) were embedded in a membrane consisting of POPC:POPG (3:2) and 8% of cholesterol, in line with the composition of the SMO-containing nanodiscs used in this study and previously^12^. The receptor was oriented using data from the OPM database^136^. On the SMO C-tail the GRK2/3 phosphorylation sites, defined here **(Extended Data Fig. 8**) and in our previous study^13^, were included in the model (phosphorylated S560, S578, S594, T597, S599, S615, S642, T644, T648, S666), while on the PKA residues S10, S197 and T338 were phosphorylated, as these residues are known to be phosphorylated in PKA-C purified from recombinant systems^137^. Complexes were solvated using TIP3P water. The net charge of the system was kept at 0.15 using NaCl ions. The generated systems were equilibrated for 50ns, with constraints applied to protein backbone and ligand heavy atoms using a timestep of 2fs. Pressure was kept at 1.0132 using the Berendsen barostat. This was followed by 3 production runs of 1µs in NVT conditions. To assess the impact of mutations on the SMO PKI-like helix and RII-like helix, we simulated the region of the SMO C-tail spanning from residue D562 to R633, using the AlphaFold 3 model of the SMO / PKA-C complex as a starting point. The protein was solvated with water and NaCl ions, and the system underwent the same NPT equilibration protocol. Subsequently each system was simulated in NVT conditions for 500 ns in 3 replicates. For experiments to examine force-induced unfolding of the SMO pCT, we have simulated the region of the SMO C-tail spanning from residue G558 to R632, using the AlphaFold 3 model of the SMO / PKA-C complex as a starting point. With constraints applied to the H558 N atom, the R632 was slowly extended, by attaching a pseudo atom which was pulled away with in the Z-plane with a constant velocity of 0.025 Å/ps. The pseudo atom was connected to R632 with an elastic bond with a spring constant of 1 kcal/mol/Å^2^. Simulations were carried out using NAMD^138^ using a timestep of 2fs. Simulations were stopped when the protein reached a fully unfolded conformation (283 Å of extension). All simulations are available in the GPCRmd platform at: https://www.gpcrmd.org/dynadb/publications/1536/.

### HDX-MS sample preparation and data acquisition

Due to the instability of the SMO / PKA-C complex during gel filtration chromatography (see main text), we used disulfide trapping to stabilize the complex. The high specificity of this crosslink (C199 of PKA-C to 637 of SMO (L637C) (**Fig. 2b**) enabled us to trap the complex without altering the biological activity. The deuterium exchange reaction was carried out by diluting 3 μL of sample at ∼15-20 μM in 57 μL deuterium exchange buffer (94.99% D_2_O, 20 mM HEPES pH 8.0, 150 mM NaCl, 0.025% Glycodiosgenin, 1 mM ATP, 10 mM MgCl_2_) for a final deuteration of 89.99%. Phosphorylated or nonphosphorylated SMO (L637C) / PKA-C complexes were prepared as described above. HDX-MS runs carried out for pSMO (phosphorylated) were performed in the presence of agonist SAG21k (1 μM) while runs with nonphosphorylated SMO were performed in the presence of inverse agonist KAAD cyclopamine (1 μM), to maintain SMO in an active or inactive conformation, respectively. To maintain SMO solubility, the buffer was also supplemented with 0.025% GDN (a non-ionic detergent that is mass spectrometry compatible). The reactions were performed for free PKA-C, free pSMO, free SMO, pSMO / PKA-C complex, and SMO / PKA-C complex. The reactions were carried out at 25° C in triplicate for deuteration times 1 min, 5 min, 10 min and 30 min following which they were quenched by the addition of 60 μL quench buffer (1.5 M guanidinium hydrochloride, 0.25 M TCEP) to bring the reaction to pH 2.5. Undeuterated control runs were also performed in a buffer without D_2_O.

100 μL of quenched samples were injected into an ACQUITY nano-UPLC HDX manager (Waters, USA) and proteolyzed in an immobilized BEH pepsin column in 0.1% formic acid running at a continuous flow rate of 100 μL/min to generate peptic peptides. The peptides were trapped in a VanGuard trap column and loaded into a C18 column and eluted in an acetonitrile gradient (8%-40%) in 0.1% formic acid following which the peptides were ionized by electrospray ionization and sprayed into a Synapt XS Quadrupole time of flight mass spectrometer where data was acquired in HDMS^E^ mode. Ion mobility settings of 600 m/s wave velocity and 197 m/s transfer wave velocity were used with collision energies of 4V and 2V used for trap and transfer respectively. Increasing high collision energy from 20-45V with a cone voltage of 20V was used to scan an m/z range of 50-2000 m/z in positive ion mode. [Glu^1^]-fibrinogen peptide B([Glu]fib) was (100 fmol/min) at a flow rate of 5 μl/min was used as a lockspray reference. The entire run time was 15 min consisting of a 3 min proteolysis and 12 min acquisition.

### HDX-MS data analysis

The murine sequences of SMO (64-674) (Uniprot P56726) and PKA-C (Uniprot P05132) were used for peptide identification in Protein Lynx Global server v3.0 (PLGS, Waters) in HDMS^E^ mode with workflow parameters set to non-specific proteolytic cleavage, variable phosphorylation at Ser/Thr/Tyr. The PLGS results were combined with the raw spectra for analysis in DynamX v3.0. The DynamX filters were minimum intensity=2000, minimum products per amino acid=0.2, minimum peptide length=5, maximum peptide length=25, maximum mass tolerance=10 ppm. Spectra were analyzed for each state by comparing the undertreated run to the time point runs and state wise comparisons were also performed. The final overall sequence coverage on SMO was 56.7% yielding 74 peptides at an amino acid redundancy of 2.17. Correspondingly the sequence coverage on PKA-C was 86.3% yielding 72 peptides at an amino acid redundancy of 2.56. Greater than 50% of all peptides in our data set showed HDX values ≥1 Da, indicating efficient proton-deuterium exchange. Back exchange was estimated to be 19.8% for the most deuterated peptide on phosphorylated SMO residues 541-546 by considering the deuteration percentage and the number of exchangeable amides from the 48hr fully deuterated data. We designated 0.5 Da as the threshold value above which differences in HDX values between experimental conditions (either positive or negative differences) are considered meaningful; this value was established based on experimental uncertainty in deuterium uptake across several proteins at a 98% confidence interval, as defined previously^139^.

Peptides displaying bimodal spectra were selected for bimodal analysis in HXexpress v with four peptides from SMO being chosen for the analysis (96-112, 142-151, 431-441 and 641-654). However, due to poor S/N ratios, peptides 96-112 and 431-441 were excluded from the analysis. Bimodal analysis was performed for peptides 142-151 and 641-654 and bimodal spectra were assessed for statistical significance using a two-parameter metric: a p-value of <0.05 and a confidence interval >95% in the regression metric. Two additional parameters were also evaluated to prevent overfitting-the delta chi metric and separation metric, as described previously^140^.

### Mapping SMO phosphorylation sites via targeted mass spectrometry

FLAG-tagged mouse SMO (residues 64-674) was purified from HEK293 cells treated with SMO modulators and/or GRK2/3 inhibitors, as described previously^13^. The purified protein extracts from different conditions were denatured and reduced in 1.7 M urea, 50 mM Tris-HCl pH 8.0, 1 mM DTT at 37°C for 30 minutes, alkylated in the dark with 3 mM iodoacetamide at room temperature for 45 minutes, and excess iodoacetamide was quenched with 3 mM DTT for 10 minutes at room temperature. For digestion, proteins were incubated with 1 μg chymotrypsin at 37°C overnight. To stop the digestion, samples were acidified with 0.5% trifluoroacetic acid (TFA). Digested samples were desalted for MS analysis using a BioPureSPE Mini 96-Well Plate (20 mg PROTO 300 C18; The Nest Group) according to standard protocols.

Digested samples were analyzed on an Orbitrap Exploris 480 MS system (Thermo Fisher Scientific) equipped with an Easy nLC 1200 ultra-high pressure liquid chromatography system (Thermo Fisher Scientific) interfaced via a Nanospray Flex nanoelectrospray source. For all analyses, samples were loaded onto a 75 μm ID C18 reverse phase column packed with 25 cm ReprosilPur 1.9 μm, 120Å particles (Dr. Maisch). Mobile phase A consisted of 0.1% FA, and mobile phase B consisted of 0.1% FA/80% ACN. Peptides were separated by an organic gradient from 2% to 28% mobile phase B over 32 minutes followed by an increase to 44%B over 19 minutes, then held at 90% B for 9 minutes at a flow rate of 300 nL/minute. Analytical columns were equilibrated with 6 μL of mobile phase A. To build a spectral library, the 4 biological replicates for each condition were pooled and acquired in a data-dependent manner. Data dependent analysis (DDA) was performed by acquiring a full MS1 scan over a m/z range of 350 to 1,250 in the Orbitrap at 120,000 resolving power (@200 m/z) with a normalized AGC target of 100%, an RF lens setting of 40%, and a maximum ion injection time set to “Auto.” Dynamic exclusion set to 30 seconds, with a 10 ppm exclusion width setting. Peptides with charge states 2 to 6 were selected for MS/MS interrogation using higher energy collisional dissociation (HCD), with a set cycle time of 1 second. MS/MS scans were analyzed in the Orbitrap using isolation width of 1.3 m/z, normalized HCD collision energy of 30%, normalized AGC of 200% at a resolving power of 15,000, and with a maximum ion injection time set to “Auto.” For all acquisitions, QCloud was used to control instrument longitudinal performance during the project^141^. All proteomic data were searched against the human UniProt database (UniProt reviewed sequences downloaded 07/2018) augmented with the sequence of the affinity tagged mouse SMO. Peptide and protein identification searches, as well as label-free quantitation were performed using the MaxQuant data analysis algorithm (version 1.6.12.0)^142^ using above described parameters. The database search results were used to generate a spectral library in Skyline (version 20.2.0.343)^143^ and to extract optimal coordinates for targeted proteomics assays (so called parallel reaction monitoring (PRM) assays). PRM measurements were performed on all 4 biological replicates per condition separately using the above-described gradient for spectral library generation but operating the Orbitrap Exploris 480 in PRM mode. Targeted MS2 spectra were acquired using the following parameters: 60 k resolution, scan range set to “Auto,” HCD with 30% NCE, RF lens setting of 50%, an AGC target set to = “Standard”, the maximum injection time set to “Dynamic,” desired minimum points across the peak set to “9”, and an isolation window of 1.2 m/z. Selected SMO peptides were targeted in 3-minute wide transition windows. The resulting data were analyzed with Skyline (version 20.2.0.343) for identification and quantification of peptides [PMID: 20147306]. MSstats was used for statistical analysis^122^. Statistical analysis of unmodified SMO and detected phosphosites was performed separately for the different digestion conditions using the statistical framework Msstats^122^. Intensities are estimated using the sample quantification function in MSstats which provides model-based estimation of phosphosite and protein abundance combining individual peptide intensities. Quantification was graphed using Prism 8 (GraphPad). Raw data and PRM transition files can be accessed, queried, and downloaded via v (https://panoramaweb.org/SMO_phospho_2024.url)^144^. Reviewer access information:

Password: I018lNrDtz?uZG

### Purification of PKA-C from native tissue

Native-source PKA-C was purified from a beef heart via IP20 chromatography. A fresh beef heart (Tooele Valley Meats; Tooele, UT) was transported on ice before the purification of PKA-C. Meat from the left ventricle of the beef heart was cut into small pieces. For 100 g of tissue, 200 mL lysis buffer (50 mM Tris-HCl pH 7.4, 50 mM NaCl, 2 mM MgCl_2_) supplemented with Pierce EDTA-free protease inhibitor tablets was added. Then the mixture was homogenized in a blender for five 1-minute cycles with cooling intervals between cycles. The homogenized material was centrifuged at 13,000 × g and 4 °C for 20 min. The supernatant was centrifuged for a second round at 38,000 × g and 4 °C for 45 min. The supernatant was filtered through an empty resin column before batch binding with IP20 resin in 50-mL conical tubes. After supplementing with 100 µM cAMP, 5 mM MgCl_2_, and 3 mM ATP, the tubes were incubated at 4 °C overnight with gentle end-to-end rotation. The IP20 resin was separated by passing the mixture through a column. The resin was washed with W1 buffer (50 mM Tris-HCl pH 7.4, 50 mM NaCl, 2 mM MgCl_2_, and 0.4 mM ATP) and W2 buffer (50 mM Tris-HCl pH 7.4, 250 mM NaCl, 2 mM MgCl_2_, and 0.4 mM ATP). Bound protein was eluted with elution buffer (50 mM Tris-HCl pH 7.4, 200 mM arginine, 50 mM NaCl, and 1 mM EDTA). The protein sample was concentrated and further purified using a Superdex 200 Increase 10/300 GL column equilibrated with storage buffer (20 mM MOPS pH 7.0, 150 mM NaCl, 2 mM β-ME). Monomeric fractions were pooled, concentrated, flash-frozen, and stored at −80 °C before use.

### BS3 crosslinking of SMO/PKA-C complex for cryoEM studies

Purified and in vitro phosphorylated mSMO (64-674) and native-source bPKA-C were mixed in binding buffer (20 mM HEPES pH 7.5, 150 mM NaCl, 0.01% GDN, 10 mM MgCl_2_, 1 mM ATP, and 1 µM SAG21k) at a 1:1.2 molar ratio and incubated at room temperature for 30 min. Fresh BS3 dissolved in binding buffer was added to a final concentration of 1 mM, and the sample was incubated at room temperature for 45 min. After incubation, the reaction was stopped by the addition of Tris-HCl pH 8.0 to a final concentration of 50 mM. Then the complex was purified using a Superdex 200 Increase 10/300 GL column equilibrated with binding buffer. The peak fractions of monomeric complex were pooled and concentrated to 10 mg/mL for cryoEM studies.

### CryoEM sample preparation, data acquisition and processing

BS3-crosslinked SMO/PKA-C complex was frozen on glow-discharged UltrAuFoil grids (Quantifoil, R 1.2/1.3, 300 mesh) at a concentration of 10 mg/mL using a Vitrobot (Thermo Fisher Scientific). Data were collected on a 300 kV Titan Krios transmission electron microscope as 40-frame movies with a total exposure of 50 e^-^/Å^2^. The raw pixel size was 1.06 Å. A total of 831 movies were collected. Movies were imported into CryoSPARC and motion-corrected with Patch Motion Correction. CTF parameters were estimated by CTFFIND4. Micrographs with CTF-fit resolution better than 3.5 Å were selected for particle picking. Template Picker was used to pick particles from the whole dataset with 2D templates generated from 100 micrographs. A total of 483,280 particles were picked for 2D classification. After 2D classification, 64,118 particles were selected for Ab-initio reconstruction. One class with 21,122 particles exhibited the expected overall features of the SMO/PKA-C complex (i.e., SMO embedded in detergent micelles and PKA-C attached to one side). However, further 3D classification and refinement failed to reveal any secondary structure. The 3D class from the first-round 3D classification was used to generate an EM map using Non-uniform Refinement.

## ACKNOWLEDGMENTS

We thank B. Graves and S. Nakielny for comments on the manuscript. P.S. was supported by an NIH T32 training grant (T32-GM122740). J.W.B. was supported by the German Research Foundation (DFG)-funded Research Training Group “multiscale clocks” (448909517/GRK 2749: Biological Clocks on Multiple Time Scales). This work was supported by NIH grants R01EY035377 (S.S.T. and B.R.M.), and R35GM133672 (B.R.M.).

## AUTHOR CONTRIBUTIONS

W.P.S., N.I., and Z.M. expressed and purified all SMO, GRK2, and PKA-C proteins (except for myr K7C PKA-C used in SPR studies), prepared phosphorylated SMO for SPR, conducted all SMO/PKA-C disulfide trapping studies, prepared phosphorylated SMO/PKA-C complexes for HDX-MS, and analyzed SMO mutants for defects in GRK2 phosphorylation. N.I. also purified SMO C-terminus for CD studies and conducted CD studies with guidance from P.S. and M.K. G.L. expressed, purified, and reconstituted SMO / PKA-C complexes for cryoEM studies, prepared cryoEM grids, and analyzed cryoEM data, with guidance from E.C. V.V. designed HDX-MS studies and acquired and analyzed HDX-MS data, with guidance from G.S.A. J.W. analyzed AlphaFold models to identify residues in key interfaces, helped design mutations for functional assays and SMO/PKA-C disulfide trapping assays, and prepared structural models for publication in PyMol. T.M.S. prepared and conducted all MD simulations, with guidance from J.S. J.W.B. carried out all SPR studies and evaluated the results with guidance from D.B. and F.W.H. J.-F.Z. conducted all GLI transcriptional reporter experiments. J.B. and G.T. expressed and purified myristylated K7C PKA-C for SPR studies. A.L. constructed stable NIH3T3 cell lines for all SMO mutants and analyzed their localization to primary cilia via confocal microscopy. I.N. expressed and purified biotinylated MSP1D1 and MSP1E3D1 scaffold proteins for SMO nanodisc reconstitutions and assisted in preparing figures. C.D.A. conducted initial SMO L637C/PKA-C disulfide trapping studies. J.X. identified SMO pS615 GRK2 phosphorylation site via mass spectrometry, with guidance from R.H. R.L.D. prepared, analyzed, and interpreted all AlphaFold 2.3.0 and most AlphaFold 3 models. R.L.D., S.S.T., and B.R.M. conceived the project. S.S.T. and B.R.M. interpreted data and provided overall project supervision. B.R.M. wrote the manuscript with assistance from W.S., N.I., and Z.M., and all authors provided feedback, comments, and revisions.

